# SAICAr-dependent and independent effects of ADSL deficiency on neurodevelopment

**DOI:** 10.1101/2020.11.23.394767

**Authors:** Ilaria Dutto, Julian Gerhards, Antonio Herrera, Alexandra Junza, Oscar Yanes, Cedric Boeckx, Martin D. Burkhalter, Sebastian Pons, Melanie Philipp, Jens Lüders, Travis H. Stracker

## Abstract

Adenylosuccinate Lyase (ADSL) functions in the *de novo* purine biosynthesis pathway. ADSL deficiency (ADSLD) causes numerous neurodevelopmental pathologies, including microcephaly and autism spectrum disorder. ADSLD patients have normal purine nucleotide levels but exhibit accumulation of the dephosphorylated ADSL substrates SAICAr and S-Ado. SAICAr was implicated in the neurotoxic effects of ADSLD, although its role remains unknown. We examined the effects of ADSL depletion in human cells and found increased DNA damage signaling, that was rescued by nucleosides, and impaired primary ciliogenesis, that was rescued by reducing SAICAr. By analyzing ADSL deficient chicken and zebrafish embryos we observed impaired neurogenesis and microcephaly, and neuroprogenitor attrition in zebrafish was rescued by reducing SAICAr. Zebrafish embryos also displayed phenotypes commonly linked to ciliopathies. Our results suggest that both reduced purine levels and SAICAr accumulation contribute to neurodevelopmental pathology in ADSLD and defective ciliogenesis may influence the ADSLD phenotypic spectrum.

## Introduction

Adenylosuccinate lyase (ADSL) is a conserved homotetrameric enzyme that catalyzes two reactions in the *de novo* purine synthesis (DNPS) pathway(*1*). Mutations in *ADSL* cause adenylosuccinate lyase deficiency (ADSLD), an autosomal recessive disorder characterized by defects in purine metabolism and heterogeneous neurological phenotypes that include lack of eye-to-eye contact, auto-aggressive behavior, speech impairment, mild psychomotor delay, transient contact defects, autism spectrum disorder, epilepsy and in some cases, microcephaly, encephalopathy, ataxia or coma vigil(*2, 3*). While the incidence of ADSLD has not been fully established, over 80 patients have been diagnosed to date and subcategorized based on their symptoms that range from premature death to milder developmental and behavioral disorders(*3*).

ADSLD can be diagnosed by detecting elevated levels of the dephosphorylated substrates of ADSL, SAICAr and S-Ado, in body fluids(*2*). As normal levels of purine nucleotides were detected in serum from ADSLD patients, the accumulation of S-Ado, and particularly SAICAr, has been proposed to play a role in the disease pathology(*2–4*). In yeast, ADSL (Ade13) loss provokes genomic instability and is lethal(*1, 5–7*). Lethality in yeast can be rescued by deletion of a number of DNPS enzymes upstream of ADSL, or the transcription factors that regulate the pathway, indicating that the accumulation of metabolic intermediates, rather than impaired DNPS, underlies toxicity(*1, 7*).

In *C. elegans,* ADSL loss caused delayed growth, infertility, reduced lifespan and locomotion defects. In some studies growth, lifespan and locomotion could be linked to the accumulation of SAICAr(*7–9*). Perfusion of rat brains with SAICAr led to cellular attrition in the hippocampus, leading to the proposition that SAICAr accumulation is neurotoxic, although the potential mechanism remains unknown(*4*). In glucose deprived cancer cells, SAICAR accumulation was shown to activate PKM2, and a number of other kinases, to promote cancer survival in glucose-limiting conditions, suggesting that purine metabolite accumulation could have distinct signaling outcomes that impact on cell behavior and fate during development(*10–12*). However, despite extensive enzymology and structural information, the underlying mechanisms by which neuropathology arises in ADSLD remain unknown.

To address the potential roles of ADSL deficiency in neurodevelopment, we systematically examined the consequences of ADSL depletion in diploid human cells and *in vivo.* We found that reduced ADSL function in human epithelial cells impaired cell cycle progression, induced DNA damage signaling and impaired primary ciliogenesis. Deletion of p53 or supplementation with nucleosides could rescue cell cycle and DNA damage signaling, respectively. In contrast, ciliogenesis defects were unaffected by p53 status or nucleoside supplementation and were dependent on SAICAr accumulation. Depletion of ADSL in chicken or zebrafish embryos impaired neurogenesis and caused developmental defects. In zebrafish this included microcephaly, which is observed in some ADSLD patients. In addition, fish embryos displayed ciliopathy related phenotypes and treatment with methotrexate, to inhibit DNPS upstream of ADSL, rescued impaired neurogenesis. Together our results indicate that ADSL depletion causes context-dependent phenotypes associated with both nucleotide depletion and metabolite production that together impact neurodevelopment.

## Results

### ADSL depletion causes p53-dependent proliferation defects

To investigate the impact of ADSL on cellular homeostasis, we depleted ADSL with a pool of four siRNAs in hTERT-immortalized human retinal epithelial cells (hTERT-RPE-1, referred to henceforth as RPE-1). Depletion of ADSL was effective with 80% depletion of the mRNA and a clear reduction in protein levels (Figures 1A, B). This was accompanied by reduced levels of AMP and GMP (Figure S1A), as well as accumulation of S-Ado (Figure 1C). We further validated the siRNA pool using one effective single siRNA (#2) (Figure S1B). As ADSL is critical for DNPS, we examined cell growth following ADSL depletion and found reduced levels of proliferation in ADSL depleted cells compared to controls (Figure 1D). ADSL depleted cells frequently lacked Ki67 expression, indicating that some cells were exiting the cell cycle, and had increased levels of p53 (Figures 1E, F). Deletion of *TP53* rescued proliferation and restored the number of Ki67-positive cells (Figure 1G, S1G), and the reduction in Ki67-positive cells could also be prevented by stable expression of an siRNA resistant allele of *ADSL* (ADSL*) (Figures 1H, S1F). Trypan blue and β-galactosidase assay indicated that there was not a detectable increase in cell death (Figure S1C) and that the Ki67 negative cells were likely quiescent and not senescent (Figure S1D). We also checked whether RPE-1 cells underwent differentiation by staining with Vimentin, a marker of undifferentiated cells(*13*), and Cytokeratin 20 (CK20), a marker of differentiation, upon ADSL depletion. We did not observe any CK20 signal or a reduction in Vimentin positive cells in the population upon ADSL silencing compared to the controls, arguing against premature differentiation (Figure S1E).

**Figure 1.**
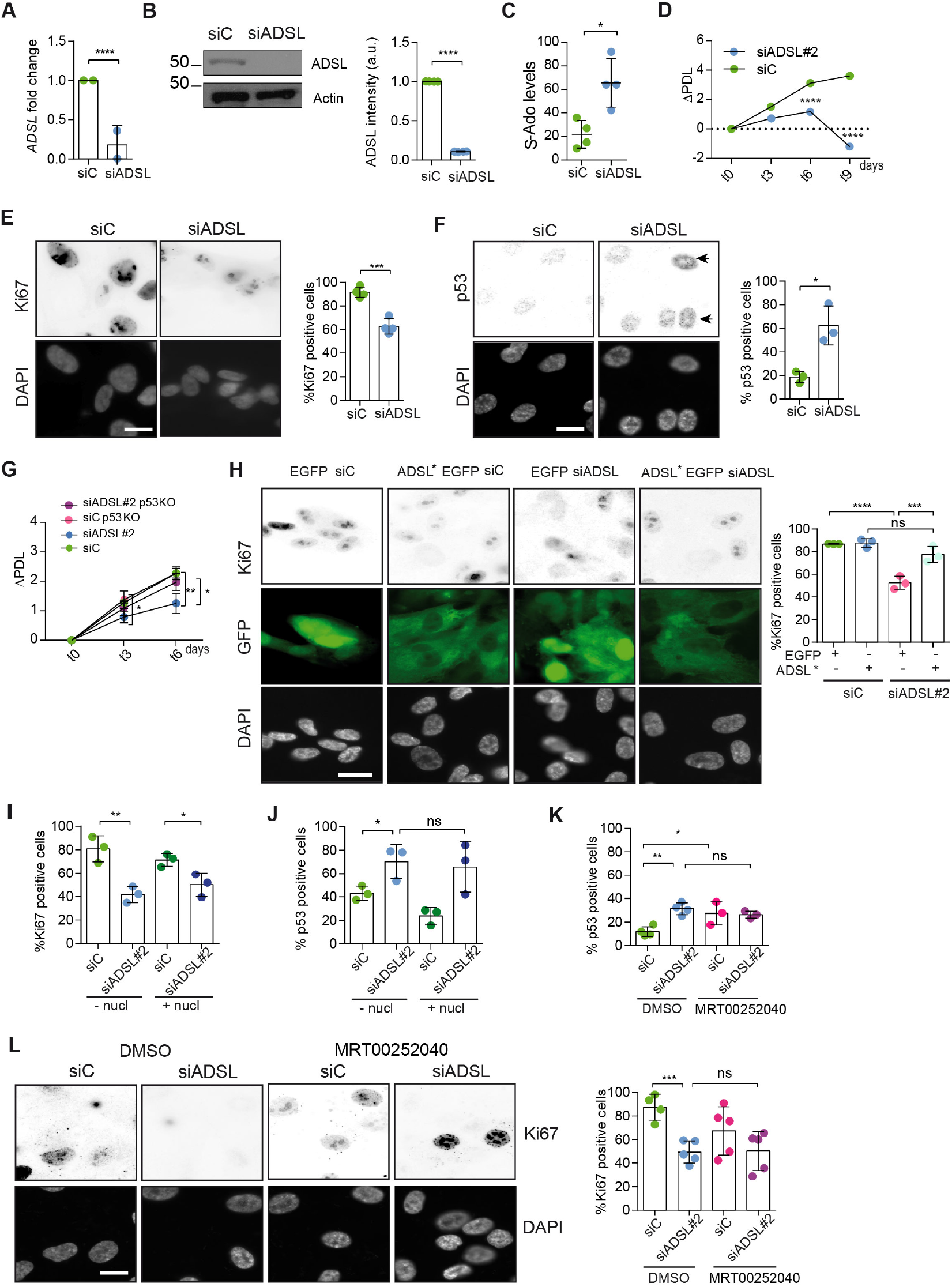
ADSL depletion causes p53-dependent proliferation defects. **A**. Reduced mRNA levels of *ADSL* confirmed by qRT-PCR experiments. hTERT-RPE-1 were silenced with smart pool RNAi for 96 hrs before harvesting. Two independent experiments in triplicate are shown in the panel (n=2 two-tailed *t*-test, ****p< 0.0001) **B**. Western blot of RPE-1 cell extracts treated as in (A). One experiment is shown as representative of three independent experiments. Actin was used as a loading control. Quantifications of ADSL intensity in four different experiments were performed by ImageJ software and normalized to actin first and then to the relative controls (n=4, two-tailed t test, ****p<0.0001). **C.** Increased S-Ado levels were detected in ADSL-depleted RPE-1 cells and compared to the controls. (n=4, two-tailed *t* test, *p<0.05). **D.** Cell proliferation rates of RPE-1 cells quantified every three days after treatment with a single control or *ADSL* siRNA in medium with serum (n=3, two-tailed *t* test ****p<0.0001). ΔPDL represent the difference in population doubling levels quantified through the formula described in materials and methods. **E**. Ki67 positive cells upon 96 hrs of silencing with control or *ADSL* smart pool siRNAs. Scale bar 10 μm. (n=4, two-tailed *t*-test, ***p<0.001). **F**. The percentage of p53 positive cells following treatment with control or ADSL smart pool siRNAs were quantified in three independent experiments (n=3, two-tailed *t*-test, *p<0.05). **G**. Cell proliferation rate in RPE-1 wt and p53 knockout KO as in (C) was counted for 6 days. (n=3, two-tailed *t* test, **p<0.01, *p<0.05). **H.** EGFP and ADSL*-EGFP stably expressing RPE-1 were transfected with a single control or ADSL siRNAs for 96 hrs and immunostained with anti-Ki67 antibody. Scale bar=20 μm. Quantification of Ki67 positive cells (n=3, one-way ANOVA test, *ns* not significant, *p<0.05, **p<0.01, ***p<0.001). **I.** Quantification of RPE-1 transfected with a single control or ADSL siRNA for 96 hrs in presence or absence of 60 μM nucleosides. Cells were fixed and immunostained with anti-Ki67 antibody. (n=3, oneway ANOVA test, *ns* not significant, **p<0.01, *p<0.05). **J**. Quantification of RPE-1 in the same conditions of (I) and immunostained with anti-p53 antibody (n=3, one-way ANOVA, *ns* not significant, *p<0.05). **K**. Quantification of p53 positive cells in *ADSL* depleted cells in the presence or absence of MRT00252040. (n=3, one-way ANOVA test, *ns* not significant, *p<0.05). **L**. Quantification of Ki67 positive cells in ADSL-depleted cells in the presence or absence of MRT00252040 (n=5, one-way ANOVA, *ns* not significant, ***p<0.001). All graphs depict means ± SD with individual values shown in circles.

To identify the cause of cell cycle exit, we supplemented cells with nucleosides, to restore purine levels, or treated with MRT00252040, a small molecule inhibitor of phosphoribosylaminoimidazole carboxylase (PAICS), to reduce elevated SAICAr levels *(14).* Supplementation of ADSL depleted RPE-1 cells with nucleosides did not prevent p53 induction or cell cycle exit (Figures 1I, J). Similarly, treatment with MRT00252040 did not influence p53 or loss of Ki67 (Figure 1K, L). This demonstrated that ADSL depletion in non-transformed human epithelial cells leads to a partial p53-dependent cell cycle exit/arrest that is not rescued by nucleoside supplementation or reduction in SAICAr levels.

**Figure S1.**
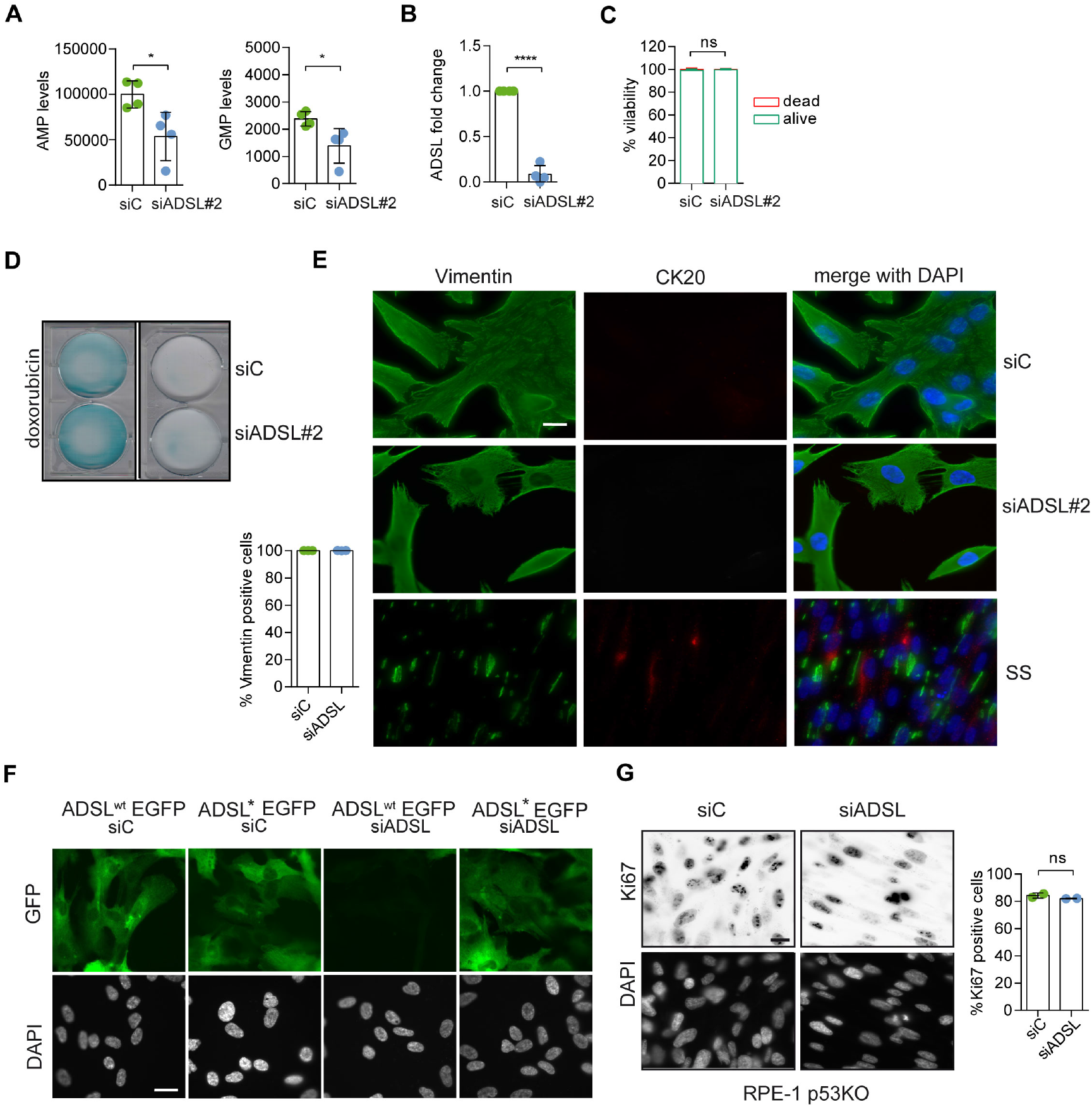
ADSL depletion reduces purine levels but does not cause senescence or promote differentiation. **A.** AMP and GMP levels in RPE-1 cells silenced with a single control or ADSL siRNA (n=4, two-tailed *t*-test, *p<0.05). **B**. qRT-PCR confirmed ADSL depletion with a single siRNA against *ADSL* (siRNA#2; n=4 in triplicate, two-tailed *t* test ****p<0.0001). **C**. Quantification of cell viability by Trypan blue in RPE-1 cells transfected with single control or ADSL siRNAs. (n=5, two-tailed *t*-test, *ns* not significant). **D.** β-galactosidase assay in RPE-1 cells upon ADSL depletion as in (B). Doxorubicin was used as a positive control. **E**. Cells were transfected with a single control or ADSL siRNA (siADSL#2) for 96 hrs, fixed and stained against Vimentin and Cytokeratin-20 (CK20). Serum starvation (SS) for 144 hrs was used as positive control for differentiation and CK20 staining. Quantification of the percentage of cells positive for Vimentin is shown. No CK20 positive cells were observed in ADSL depleted cells. Scale bar=20 μm. **F**. Control of ADSL depletion and siRNA resistant mutant expression 96 hrs post ADSL depletion. Scale bar=20 μm. **G**. Ki67 staining in RPE-1 p53 KO upon silencing with a single control or ADSL siRNA (siRNA#2; n=2, two-tailed *t*-test, *ns* not significant). Scale bar=20 μm. All graphs depict means ± SD with individual values shown in circles.

### ADSL depletion causes elevated DNA damage signaling

Reduced levels of purine nucleotides in ADSL-depleted cells may cause replication stress and DNA damage, which could trigger p53 activation (*15–17*). We observed an increased number of cells with more than five 53BP1 foci per cell, indicative of DNA double strand break accumulation (Figures 2A). 53BP1 foci were reduced by treatment with a small molecule inhibitor for ATM (Figure 2B), indicating an active DNA damage response. Supplementation of cells with nucleosides suppressed the appearance of DNA double strand breaks detected by 53BP1 and γH2AX staining (Figures 2C, D). In contrast, the PAICS inhibitor MRT00252040 did not rescue DNA double strand breaks (Figure 2E). These data indicate that ADSL depletion in cultured cells induces mild levels of DNA damage signaling that can be suppressed by nucleoside supplementation and that p53-dependent cell cycle exit is not solely a consequence of DNA damage signaling or purine metabolite accumulation.

**Figure 2.**
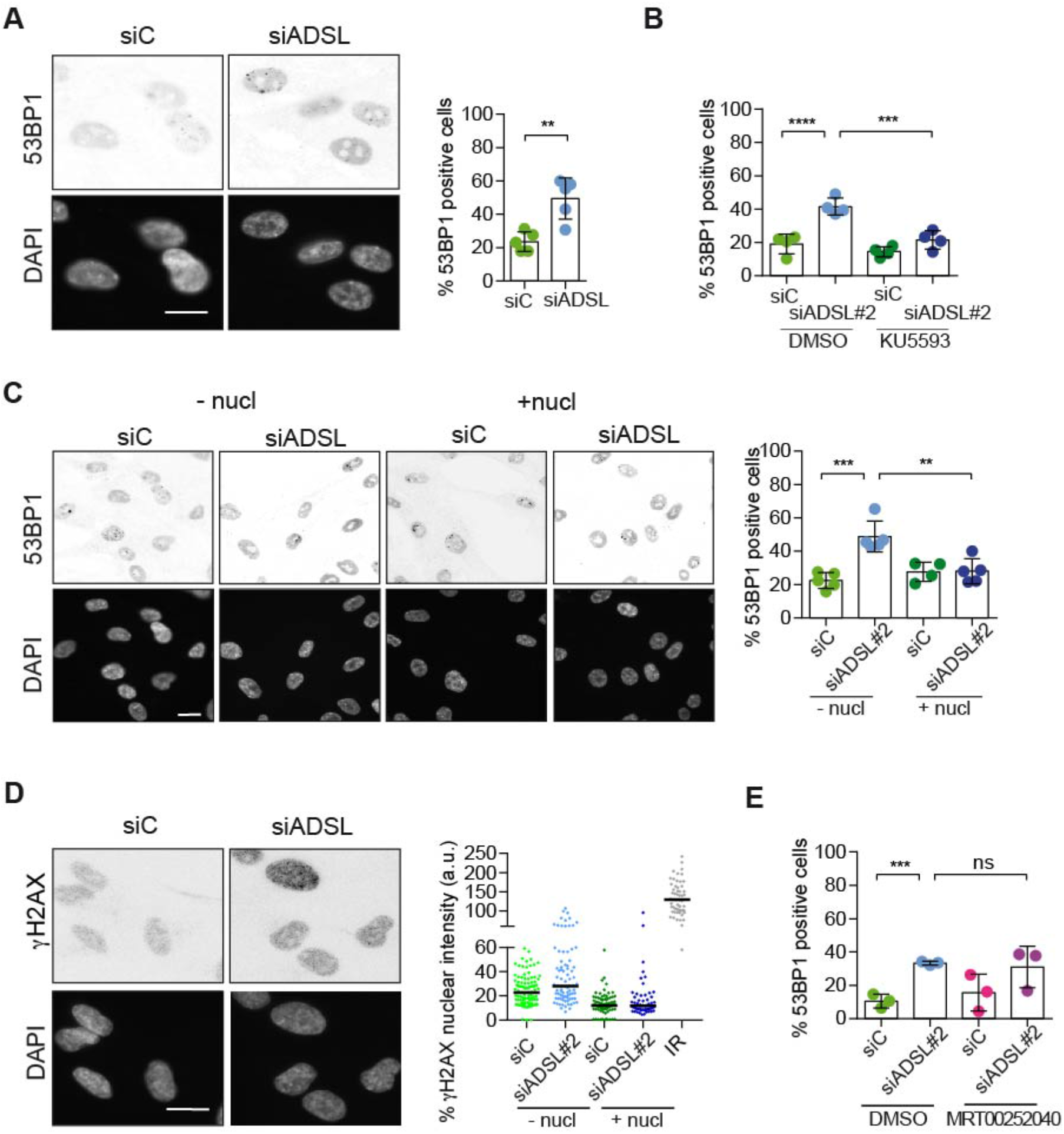
ADSL depletion caused elevated DNA damage signaling. **A.** RPE-1 were silenced for 96 hrs with a smart pool of ADSL siRNAs, fixed and immunostained with anti-53BP1 antibody. Scale bar=10 μm. Quantification of positive cells that have more than 5 foci per cell (n=5, two-tailed *t*-test, **p<0.01). **B.** RPE-1 were silenced with a single control or ADSL siRNA with or without 5 mM ATM inhibitor (KU5593) (n=4, one-way ANOVA test, ****p<0.0001, ***p<0.001). **C.** Cells were silenced for 96 hrs, treated or not with 60 μM nucleosides and stained for 53BP1. Scale bar=10 μm. (n=5, one-way ANOVA test, ***p<0.001, **p<0.01). **D.** RPE-1 treated as in (A) were fixed and stained for γH2AX (H2AX phosphorylated on Ser-139). Scale bar=10 μm. 5 Gy X-ray irradiation (IR) was used as positive control. Quantification of one representative experiment of two that showed similar results is shown; median is indicated in black. After normalization to the average of the control (siC), one-tailed *t* test was used for statistical analysis of *n*=3 independent experiments: *p<0.05was observed for siADSL (to siC), and for siADSL relative to siADSL+nucl. There is no statistical difference between siC and siC+nucl. **E.** RPE-1 were silenced in the presence or absence of 4 μM MRT00252040, fixed and stained for 53BP1 (n=4, one-way ANOVA test, *ns* not significant, ***p<0.001). All bar graphs show means ± SD with individual values in circles.

### ADSL depletion impairs neurogenesis in the developing chicken neural tube

Given the effects of ADSL depletion on cell growth and proliferation, we sought to examine the consequences of its loss *in vivo.* To this end, we used the chicken embryo system to examine the influence of ADSL depletion on nervous system development. We electroporated one side of the neural tube with plasmid expressing *GFP* as a transfection marker in combination with either control or ADSL shRNA vectors. After confirming efficient ADSL depletion (Figure 3A) we evaluated neurogenesis by staining with markers for proliferating neural progenitors (SOX2 positive) and post-mitotic neurons (ELAVL3/4 positive). We found that in the ADSL depleted side, both cell populations were reduced when compared to the non-transfected side (Figure 3B) and that the size of the tissue was smaller, suggesting reduced growth and/or increased cell death. Staining for the apoptotic marker Cleaved-Caspase-3 revealed no notable differences, suggesting that this was not due to increased cell death (Figure S2A).

**Figure 3.**
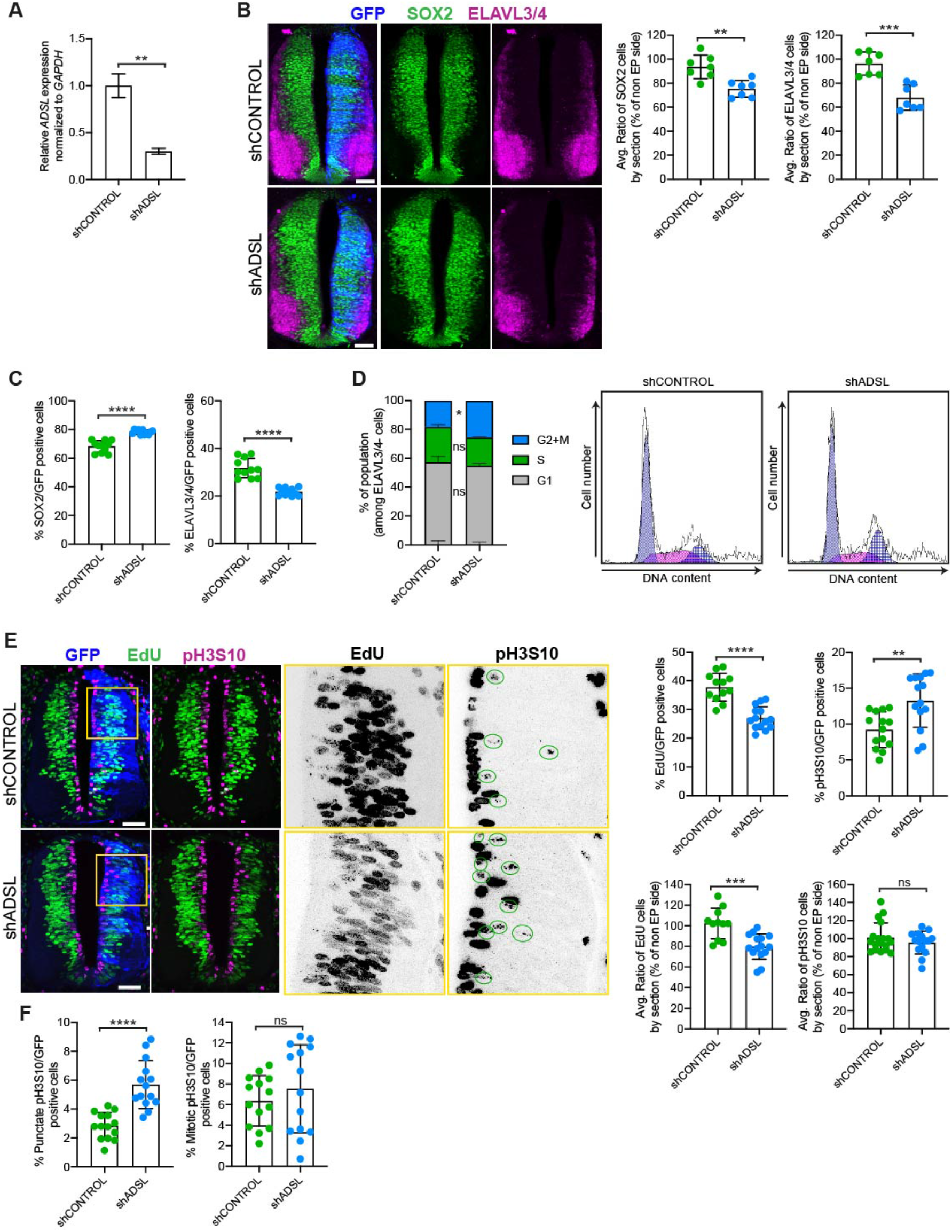
ADSL depletion causes neurodevelopmental delay in the chicken neural tube. **A**. mRNA levels of *ADSL* and *GAPDH* were measured by qRT-PCR in chicken embryonic fibroblasts (CEFs) transfected for 24 hrs with shCONTROL and shADSL to confirm knockdown efficiency (n=3, two-tailed *t*-test, **p<0.01). **B**. Transverse sections of HH12 chicken neural tubes 48 hrs post electroporation (hpe) with shCONTROL and shADSL plasmids and stained with antibodies against SOX2 (green) and ELAVL3/4 (magenta). Transfection was detected by GFP (blue). Scale bar=50 μm. Average ratio of neural stem cells (NSCs, SOX2+) 48 hpe with shCONTROL and shADSL obtained by comparing the mean number of SOX2+ cells on the electroporated and non electroporated side (n=7 embryos, two-tailed t-test, **p<0.01). Average ratio of cells differentiated into neurons (ELAVL3/4) at 48 hpe with shCONTROL and shADSL obtained by comparing the mean number of ELAVL3/4 positive cells on the electroporated and the non electroporated side (n=7 embryos, two-tailed *t*-test, ***p<0.001). **C.** Percentage of electroporated cells indentifed as NSCs (SOX2) or neurons (ELAVL3/4) 48hpe with shCONTROL and shADSL (n=11 embryos, two-tailed *t*-test, ****p<0.0001). **D**. The cell cycle profiles of NSCs (GFP+/ELAVL3/4-) obtained by FACS 48 hpe with shCONTROL and shADSL into HH12 chicken neural tubes. The mean of two independent experiments is shown in the left panel. 6-8 embryos per condition were used for each experiment. Two-tailed *t*-test was used for statistical analysis of *n=2* independent experiments, *ns* not significant, *p<0.05 Cell cycle profiles of a representative experiment are shown in the right panels. **E.** Transverse sections of HH12 chicken neural tubes 48 hpe with shCONTROL and shADSL plasmids, and stained with EdU (green) and an antibody against pH3S10 (magenta). Transfection was detected by GFP (blue). Scale bar=50 μm. Areas indicated in yellow are amplified in the right panels showing separated channels in black. Green circles in pH3S10 amplification show punctate pH3S10 positive cells. Percentage of transfected cells indentifed as EdU 48 hpe with shCONTROL and shADSL (n=12 embryos (shCONTROL) and 14 embryos (shADSL), two-tailed t-test, ****p<0.0001). Percentage of pH3S10 among the GFP+ cell population 48 hpe with shCONTROL and shADSL (n=14 embryos, two-tailed t-test, *ns* not significant, **p<0.01, ****p<0.0001). Average ratio of EdU and pH3S10 positive cells 48 hpe of shCONTROL and shADSL plasmids, obtained by comparing the mean number of EdU cells on the electroporated and the non electroporated side (EdU: n=11 embryos (shCONTROL), 15 embryos (shADSL), two-tailed t-test, ***p<0.001; pH3S10: n=18 embryos (shCONTROL), 15 embryos (shADSL), two-tailed t-test, *ns* not significant). **F.** Percentage of punctate pH3S10 (G2 phase) and mitotic pH3S10 (M phase) among the GFP+ cell population 48 hpe of shCONTROL and shADSL plasmids (n=14 embryos, two-tailed *t*-test, *ns* not significant, ****p<0.0001). Bar graphs show means ± SD.

We then analyzed SOX2 and ELAVL3/4 staining only within the GFP-positive transfected cells and found that ADSL depletion increased the percentage of SOX2-positive progenitors relative to ELAVL3/4 positive neurons (Figure 3C, S2B). This suggested that reduced tissue growth was not due to premature differentation but possibly due to a proliferation defect in the progenitor population. To study cell cycle progression in neural stem cells we performed FACS analysis of GFP-positive, ELAVL3/4 negative cells following electroporation of control and ADSL shRNA. We found that there was a slight increase in the G2/M population after ADSL depletion (Figure 3D). Further analysis of stained tissue sections showed that ADSL depletion caused a reduction in the fraction of cells that incoporated EdU and an increase in the fraction of cells positive for the G2/M marker phosphorylated Histone H3-Ser10 (pH3S10) (Figure 3E). We separated the pH3S10 positive cells into two populations; G2 cells, identified by punctate pH3S10 staining, and mitotic cells, dispaying broadly distributed pH3S10 staining. This revealed that only the G2 fraction of cells was increased by ADSL depletion, indicating that ADSL depletion caused a specific delay in G2 phase, rather than during mitosis (Figure 3F). Together our data indicate that ADSL depletion leads to a mild induction of DNA damage signaling and impaired cell cycle progression. *In vivo,* this manifests as reduced cellularity in the developing brain, without a clear induction of cell death or senescence.

**Figure S2.**
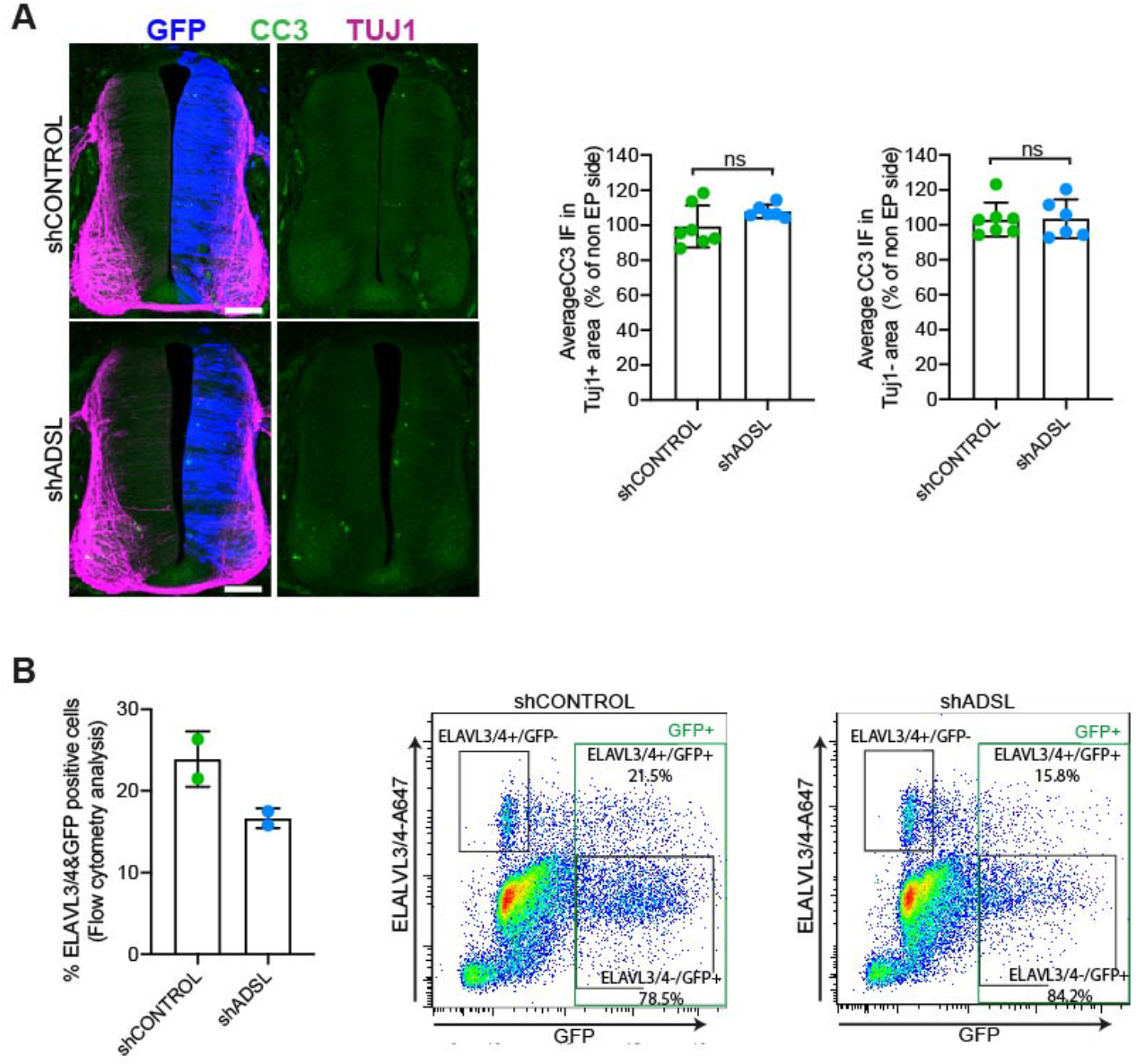
Lack of cell death or increased differentiation in developing ADSL-depleted chicken neural tubes. **A.** Representative transverse neural tube sections and quantification of mean Cleaved-Caspase-3 (CC3; green) immunofluorescence intensity obtained by comparing mean CC3 intensity on TUJ1-or TUJ1+ area (magenta) on the electroporated side (GFP area, blue) with the respective area on the non electropated side after 48 hpe with shCONTROL and shADSL (n=7 embryos (shCONTROL) and 6 embryos (shADSL), two-tailed t-test, *ns* not significant). Scale bar=50 μm. **B.** Rate of differentiation was analyzed by FACS after 48 hpe into HH12 neural tubes with shCONTROL and shADSL. 6-8 embryos per condition were used for each experiment. The mean of two independent experiments are shown in the left panel. Dot plots in right panels representing ELAVL3/4 intensity versus GFP intensity of a representative experiment.

### SAICAr-dependent ciliogenesis defects following ADSL depletion

As there are non-cycling cells in the brain and since ADSL depletion caused cell cycle exit in RPE-1 cells, conditions frequently accompanied by ciliogenesis, we tested the ability of control and ADSL depleted RPE-1 cells to assemble cilia. Following treatment with siRNA, cells were serum-starved for 48 hours and analyzed by immunofluorescence microscopy. Ki67 staining confirmed that most of the cells in both conditions exited the cell cycle (Figure 4A). We next examined ciliogenesis by staining with antibodies for ARL13B to label cilia and pericentrin (PCNT) a marker of centrosomes. Fewer cells treated with the ADSL siRNA pool had cilia and the cilia that were present were shorter when compared to controls (Figure 4B). We also observed shorter cilia upon depletion with single siRNAs (Figure S3A). To exclude the possibility that ciliogenesis was simply delayed, we quantified the number of ciliated cells 72 hrs after serum starvation and observed a similar defect (Figure S3B). Defective ciliogenesis was rescued by expression of an siRNA resistant cDNA (ADSL*) but not by nucleoside supplementation (Figures 4C, D). Inhibition of the DNPS pathway with methotrexate (MTX) had no effect on ciliogenesis in control cells (Figure S3C) but rescued both the number of ciliated cells and cilia length when ADSL was depleted (Figure S3D).

**Figure 4.**
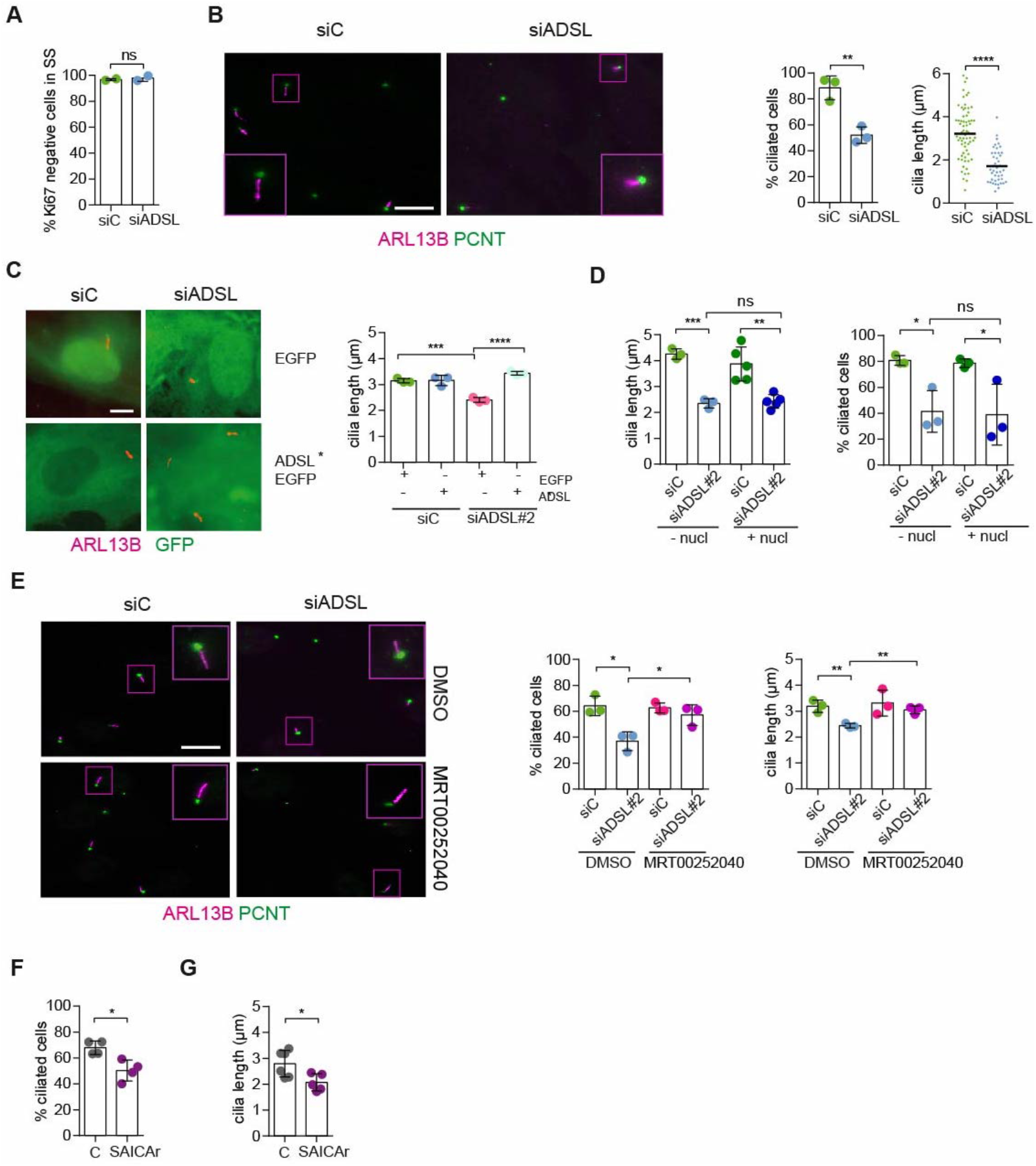
SAICAr-dependent ciliogenesis defects following ADSL depletion. **A.** RPE-1 were transfected with control and ADSL smart pool siRNAs. After 96 hrs cells were serum starved for 48 hrs to induce ciliogenesis followed by staining against Ki67 (n=2, two-tailed *t*-test, *ns* not significant). **B.** Ciliated cells silenced as in **(A)** were stained for ARL13B (magenta) and PCNT (green). Scale bar=10 μm. Magenta squares show enlargements of the areas. Graphs show quantification of ciliated cells and cilia length (line indicates median) (n=3, two-tailed *t*-test, ****p<0.0001, **p<0.01). **C.** EGFP and ADSL*-EGFP stably expressing RPE-1 were silenced for 96 hrs with control and a single ADSL siRNA, serum starved for 48 hrs, fixed and stained for ARL13B (red). Scale bar=5 μm. Graphs summarizes three experiments (one-way ANOVA, *ns* not significant, ***p<0.001, **p<0.01, *p<0.05). **D.** RPE-1 silenced with a single *ADSL* siRNA (siADSL#2) for 96 hrs in the absence or presence of 60 μM nucleosides. Cilia frequency and cilia length were quantified (mean ± SD of n=3 siC and siADSL, n=5 for siC and siADSL with nucleosides, one-way ANOVA test, *ns* not significant, ***p<0.001, **p<0.01, *p<0.05). **E.** RPE-1 cells were ADSL-depleted, treated or not with MRT00252040 and serum starved, and then immunostained for ARL13B (magenta) and PCNT (green). Scale bar=10 μm (n=3, one-way ANOVA, *p<0.05). Cilia frequency and cilia length were quantified (one-way ANOVA test, **p<0.01). **F.** Quantification of the cilia frequency in control and SAICAR-treated cells (n=4, two-tailed *t* test, *p<0.05). **G.** Cilia length measurement of cells treated as in **(F)** (n=5, two-tailed *t* test, *p<0.05).

Since ciliogenesis was rescued by MTX, which inhibits multiple steps in the DNPS pathway up and downstream of ADSL, but not by nucleoside supplementation, we next examined the potential role of SAICAr accumulation. Following ADSL depletion, we treated cells with the inhibitor MRT00252040 to specifically inhibit PAICS(*14*). This rescued ciliogenesis, as number and length of cilia were similar in control and ADSL-depleted cells (Figure 4E). Consistent with this, treatment of cells with SAICAr recapitulated the ciliogenesis defect observed in ADSL depleted cells (Figures 4F, G). To exclude indirect effects on ciliogenesis by DNA damage and resulting p53 activation, we repeated the experiment in p53 KO cells. While the overall percentage of ciliated cells was slightly lower in p53 KO cells, depletion of ADSL recapitulated the result obtained in RPE-1 wt, a reduction in ciliated cells compared to controls (Figure S3E). We concluded that SAICAr accumulation caused by ADSL depletion impaired the generation of primary cilia.

**Figure S3.**
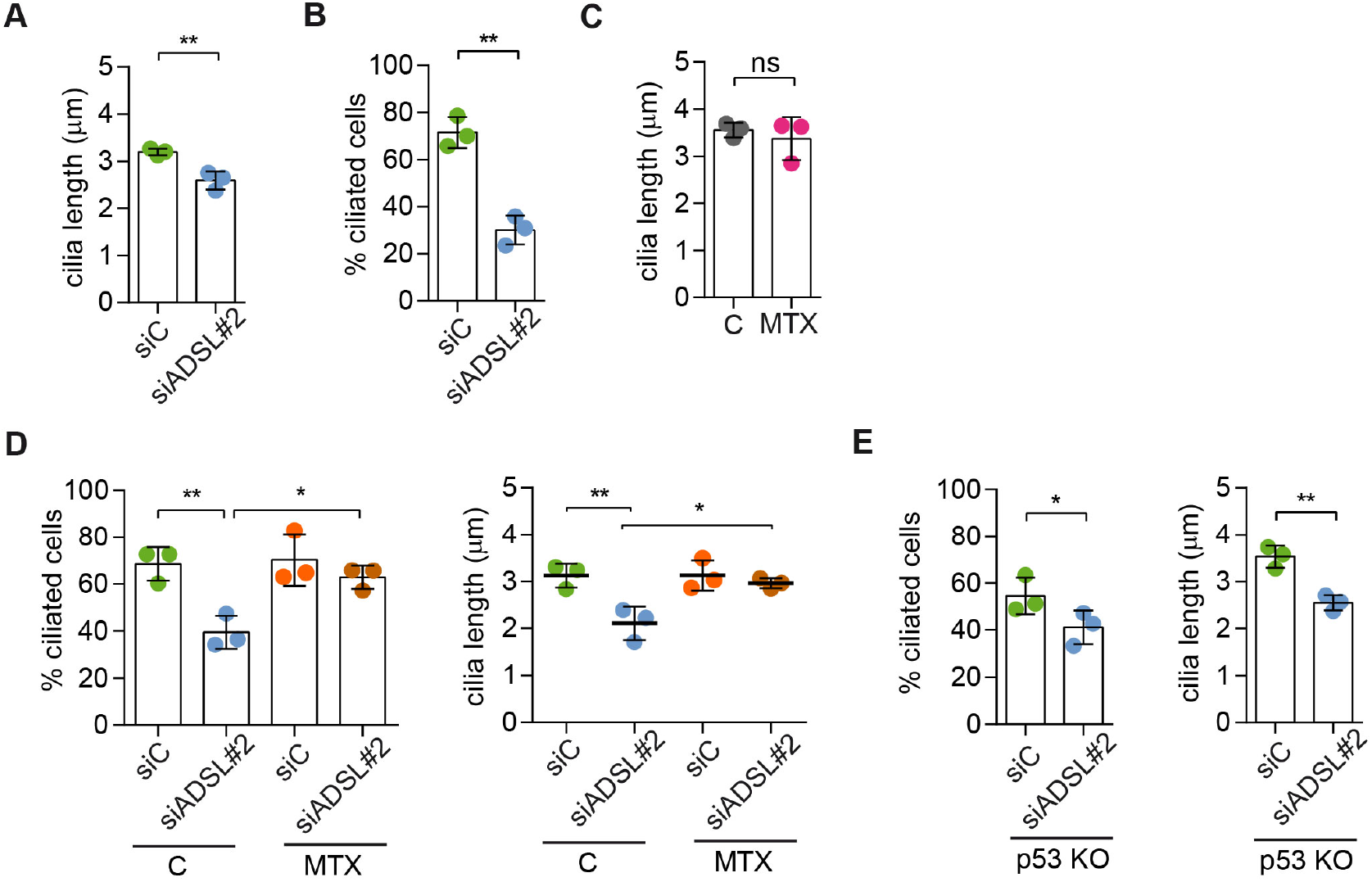
ADSL-depletion impairs ciliogenesis that can be rescued by MTX treatment. **A.** Quantification of cilia length in RPE-1 cells transfected with a single control or ADSL siRNA (n=3, two-tailed *t*-test, **p<0.01). **B.** Quantification of cilia frequency in RPE-1 silenced as in (A) and serum starved for 72 hrs. (n=3, two-tailed *t*-test **p<0.01). **C.** Quantification of cilia length in RPE-1 treated for 24 hrs with 5 μM MTX (DHFR inhibitor) (n=3, two-tailed *t*-test, *ns* not significant). **D**. Quantification of cilia frequency and cilia length in RPE-1 transfected with single control and ADSL siRNAs treated or not with 5 μM MTX for 24 hrs before serum starvation (n=3, one-way ANOVA, **p<0.01, *p<0.05). **F**. Quantification of the number of ciliated cells and cilia length in RPE-1 p53KO upon ADSL depletion (n=3, two-tailed *t*-test, **p<0.01, *p<0.05).

### SAICAr impairs CP110 removal

To understand the origin of the ciliogenesis defect, we examined centriole configurations, since mother centrioles, after conversion to basal bodies, template formation of the primary cilium. Centrosomes in ADSL depleted cells had normal levels of PCNT and normal number of centrioles (Figure 5A and S4A). However, we found that the removal of CP110 from the mother centriole, a key step in early ciliogenesis, was impaired in serum-starved, ADSL depleted cells. Compared to controls a larger number of ADSL depleted cells contained centrosomes with 2 CP110 foci (Figure 5B). This could be phenocopied by administration of SAICAr and it was rescued by PAICS inhibition (Figures 5B, C). To determine if the retention of CP110 could underlie the phenotype, we co-depleted CP110 with ADSL using three different siRNAs. All three siRNAs silenced CP110 as verified by western blot (Figure S4D) and partially depleted CP110 at centrioles (Figure S4B). In non-serum-starved conditions CP110 siRNA treated cells had fewer than the two centriolar CP110 foci typically observed in control cells (Figure S4B). Remaining centriolar signal was associated with daughter centrioles (distal to the base of the cilium in ciliated cells; Figure S4C). Co-depletion of CP110 with ADSL rescued the ciliogenesis defect (Figures 5D). These data demonstrated a SAICAr-dependent impairment of primary ciliogenesis that can be rescued by CP110 depletion or inhibition of PAICS, but not by restoration of purine levels.

**Figure 5.**
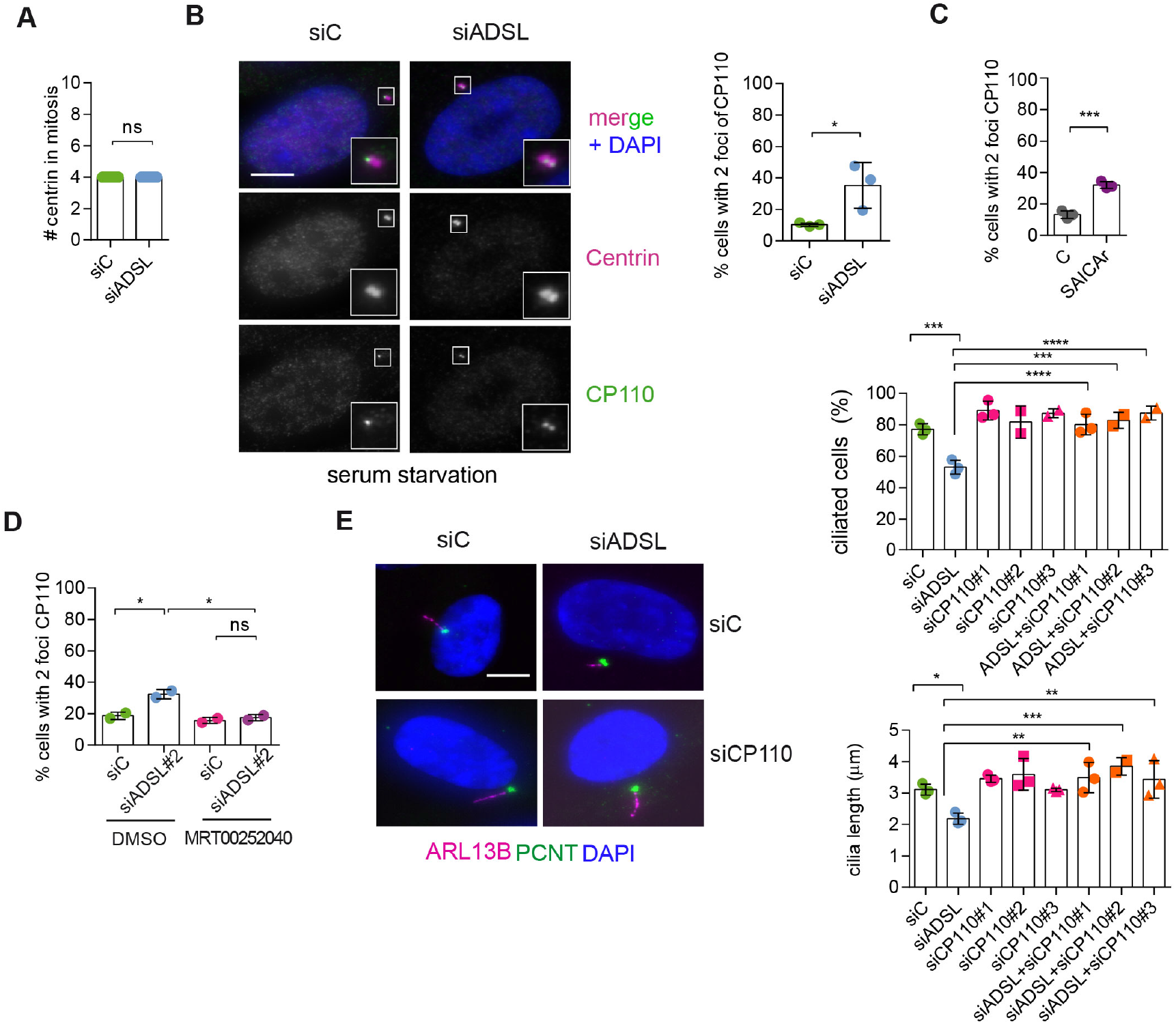
SAICAr impairs CP110 removal. **A.** Quantification of the number of Centrin foci present in mitotic RPE-1 cells transfected with control or ADSL smart pool siRNAs for 96 hrs (n=2, two-tailed *t* test, *ns* not significant). **B.** ADSL silenced cells and controls were stained for Centrin (magenta) and CP110 (green). Nuclei are shown by DAPI (blue). Graph depicts the number of ciliated cells with two CP110 foci per centrosome. (n=3, one-way ANOVA, *ns* not significant, *p<0.05). **C.** Cells mock or treated with SAICAr were processed and analyzed as described in panel (**B**) (n=3, two-tailed *t*-test, ***p<0.001). **D**. RPE-1 depleted with ADSL and control siRNAs were treated with vehicle or MRT00252040 and stained as in (**B,C**). Graph depicts the percentage of cells presenting 2 CP110 foci per centrosome (n=2; one way ANOVA, *p<0.05). **E.** RPE-1 depleted with ADSL and/or CP110 (silenced for 24 hrs with three different siRNAs) were serum starved for 48 hrs, fixed and stained for ARL13B (magenta) and PCNT (green). Graphs show number of ciliated cells (n=3 for siC, siADSL, siCP110#1, siADSL+siCP110#1; n=2 for siCP110#2, siCP110#3, siADSL+siCP110#2 and siADSL+siCP110#3, one-way ANOVA, ****p<0.0001, ***p<0.001) and cilia length (n=3, one-way ANOVA ***p<0.001, **p<0.01, *p<0.05). All graphs show means ± SD with individual values shown in circles.

**Figure S4.**
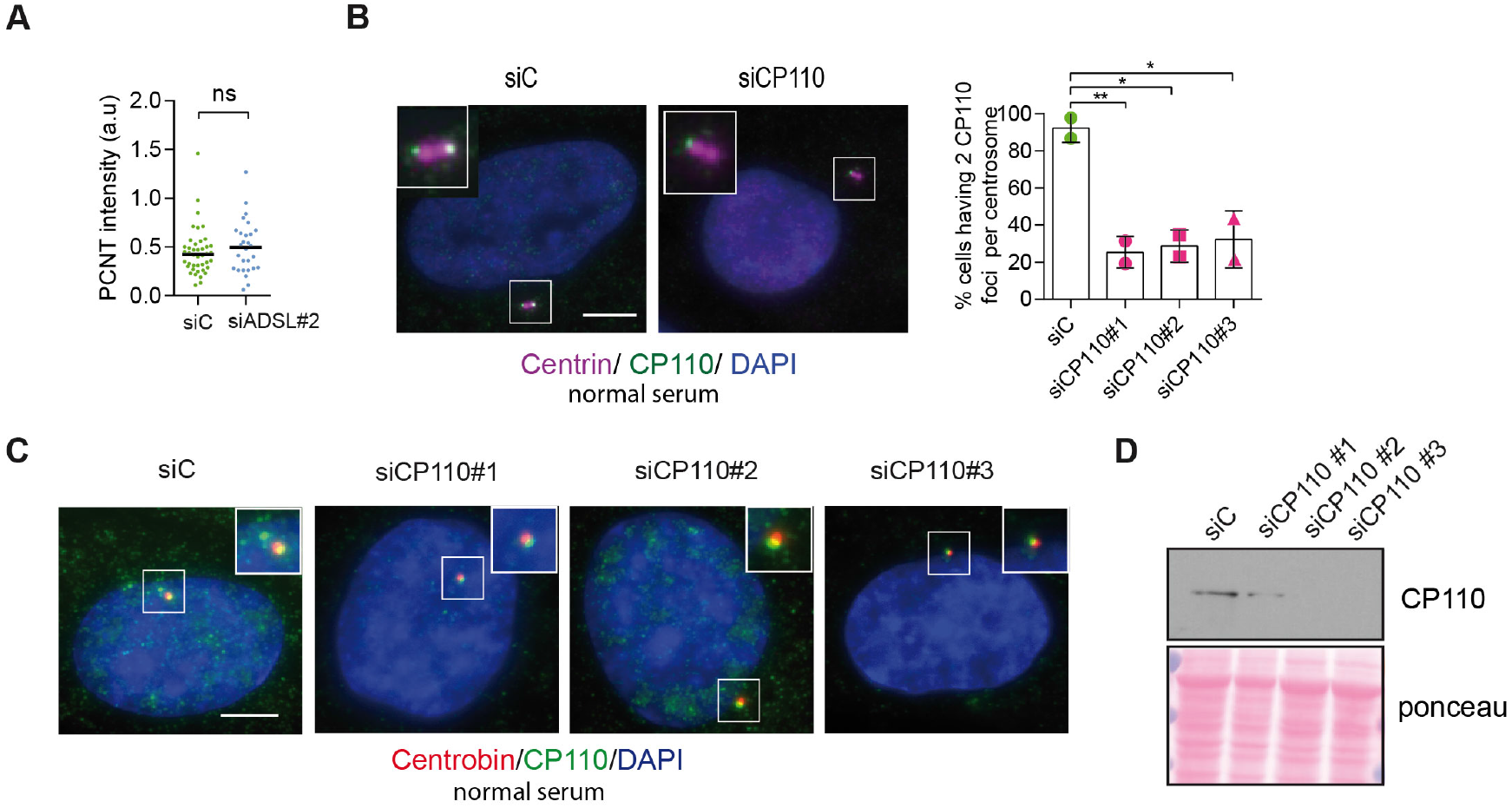
Analysis of Pericentrin accumulation and CP110 depletion. **A.** Quantification of PCNT intensity upon ADSL depletion with a single siRNA#2 (n=2, twotailed *t* test applied, *ns* not significant; median is shown). **B.** CP110 presence in both centrioles (stained by anti-Centrin antibody, in magenta) upon 24 hrs of CP110 depletion in normal serum (10%) conditions. Quantification of centrioles presenting two foci per centrosome in control and in CP110 depleted cells with three different siRNAs (n=3, oneway ANOVA test **p<0.01, *p<0.05). Scale bar=5μm. **C.** CP110 foci (in green) colocalizing with Centrobin (marker of daughter centriole, in red) upon CP110 depletion for 24 hrs in normal serum (10%). Three different siRNAs were used. Scale bar=5μm**. D**. Western blot to confirm CP110 depletion after 24 hrs of silencing.

### Depletion of Adsl in zebrafish results in developmental defects

To test whether ADSL deficiency caused ciliary defects *in vivo,* we employed a zebrafish model. As CRISPR/Cas9-mediated gene knockout did not yield viable mutants, we used two different antisense morpholino oligonucleotides (MO) to deplete Adsl in zebrafish embryos. *Adsl* is ubiquitously expressed at early embryonic stages and, by the 18-somite stage, highly expressed in several areas of the developing brain, including the midbrain and mesencephalon (Figure S5A-L). Antibody staining demonstrated expression of Adsl in neurons, which was abolished upon injection of either MO (Figure S6). Examination of embryo morphology 48 hrs post fertilization (hpf) revealed pericardial edema, kinked tail, hydrocephalus and pinhead (microcephaly) phenotypes (Figure 6A-E). Defects in head size, which are consistent with the clinical presentation of ADSLD patients, were further corroborated by staining for skull formation that is coordinated with brain development. Alcian blue staining showed that nearly 50% of the Adsl depleted embryos exhibited weak or absent staining (Figure 6F). Defects in skull formation could be largely rescued by zebrafish *Adsl* or human *ADSL* expression but not expression of a human ADSL R426H mutant, the most frequently observed ADSLD mutation (Figure 6F). Examination of DNA damage signaling in the developing neural tube revealed an increase in γH2AX positive cells. Similar to what was observed in RPE-1 cells, treatment with nucleosides suppressed DNA damage signaling (Figure 6G). These data demonstrated that Adsl depletion strongly impaired normal zebrafish development, leading to DNA damage that could be suppressed with nucleoside supplementation and several phenotypes consistent with ciliary defects.

**Figure 6.**
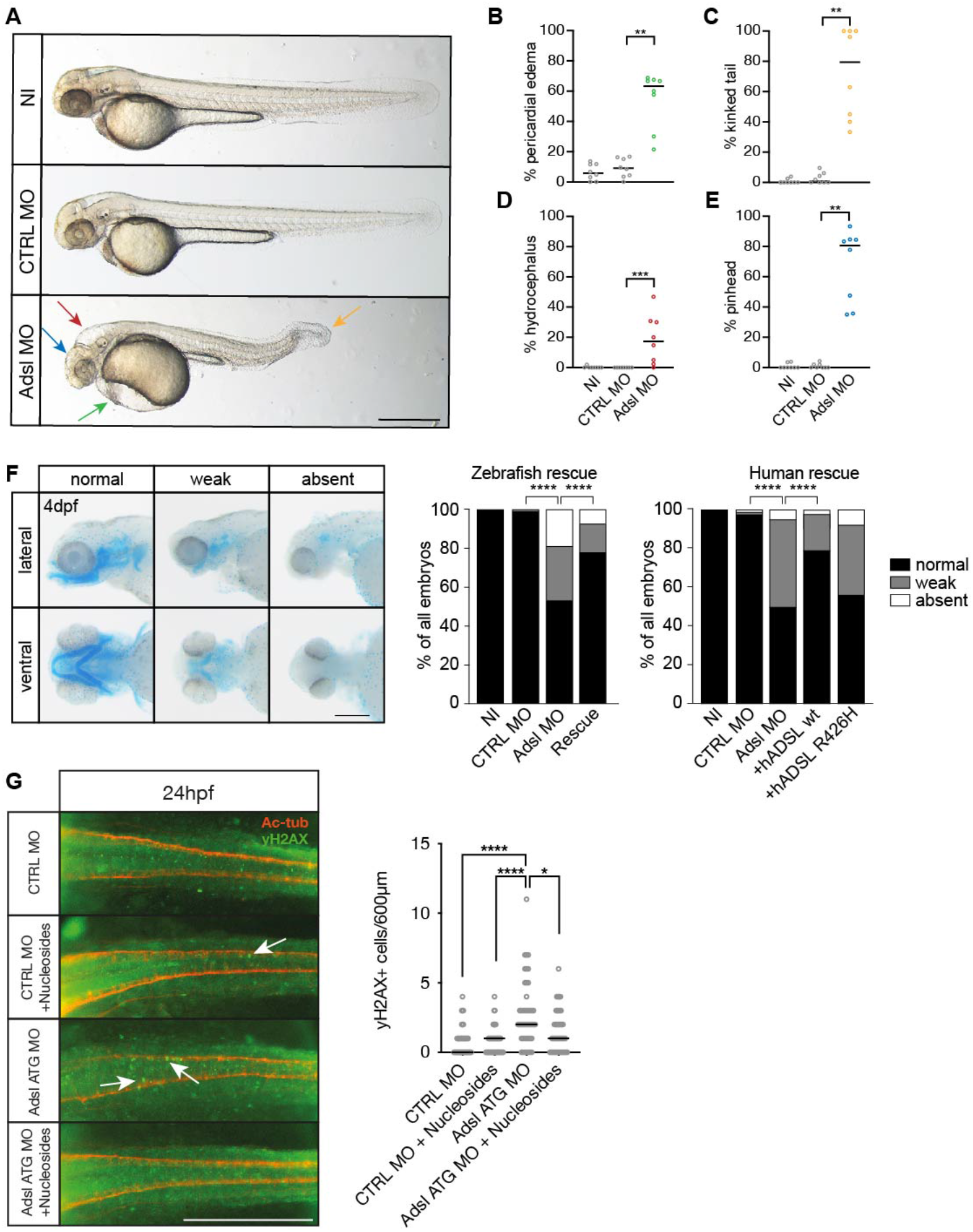
Depletion of Adsl in zebrafish causes developmental phenotypes and DNA damage signaling. **A.** Live images of 48 hpf zebrafish embryos showing pericardial edema (green arrow), kinked tail (yellow arrow), hydrocephalus (red arrow) and pinhead (blue arrow). NI (uninjected controls), CTRL MO (embryos injected with a standard control MO), Adsl ATG MO (injected with a translation blocking MO against Adsl). Scale bar=500μm. **B-E.** Quantification of the percentage of embryos developing the indicated phenotypes. For (B-E) Each circle indicates one experiment. Data from 8 experiments with 311 embryos (NI), 275 (CTRL MO), 227 (Adsl ATG MO) is shown. Kruskal-Wallis test with Dunn’s multiple comparison. Dashes show median. **p=0.0042 (pericardial edema), **p=0.0032 (kinked tail), **p=0.0011 (pinhead), ***p=0.0005 (hydrocephalus). **F.** Adsl depleted zebrafish display skull formation defects. Cartilage staining of zebrafish embryos (4 days post fertilization, dpf) with Alcian blue. Embryos were classified according to the severity of their phenotype in normal staining, weak staining or absent cartilage. Lateral and ventral view. Cartilage formation could be rescued by co-injection of capped mRNA encoding zebrafish Adsl. 6-8 experiments with a total of 178 embryos (NI), 133 (CTRL MO), 169 (Adsl ATG MO), 123 (Rescue). Injection of mRNA encoding human wt ADSL, but not the R426H ADSLD variant, restores cartilage formation in embryos. 4 experiments with a total of 116 embryos (NI), 81 (CTRL MO), 80 (Adsl ATG MO), 91 (+ *hADSL wt)* and 89 (+ *hADSL R426H).* Two-tailed Fisher’s exact test; **** p<0.0001. Scale bar=200μm. **G.** Immunofluorescence staining of the neural tube (dorsal view) of control and Adsl depleted embryos 24 hpf for γH2AX (green) and Acetylated-tubulin (Ac-tub: red). Treatment with 60mM nucleosides was carried out in indicated samples. Experiments with 45 embryos per treatment are shown, dashes indicate median. Data were analyzed by using Kruskal-Wallis test with Dunn’s correction. *p<0.05, ****p<0.0001. Scale bar=300μm. Unless indicated, comparisons are not significant.

**Supplemental Figure S5.**
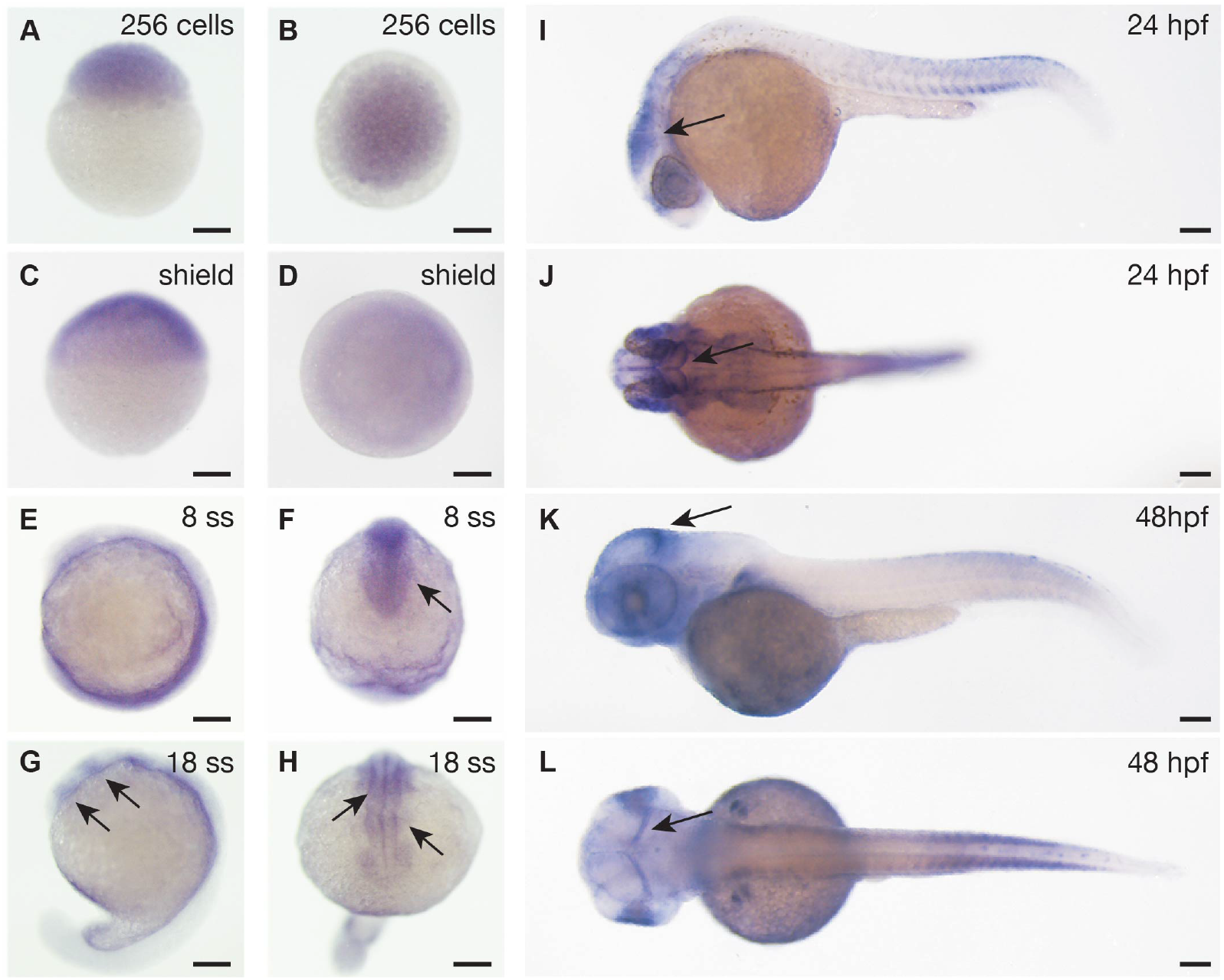
*Adsl* expression in zebrafish development. Whole mount *in situ* hybridization for detection of *adsl* expression during zebrafish development. All scale bars=100 μm. **A-D**. *adsl* is ubiquitously expressed. **E-F.** *adsl* is expressed in the anterior part of the embryo including the optic primordium (arrow in F). **G-H.** At 18 somite stage (ss) *adsl* is expressed in the developing midbrain and hindbrain (arrows). **I-J**. *adsl* is expressed in several areas of the brain including the mesencephalon (arrows). **K-L.** *adsl* is expressed in several areas of the brain including the midbrain hindbrain boundary (arrows). **D, F, H, J, L**. dorsal views.

**Supplemental Figure S6.**
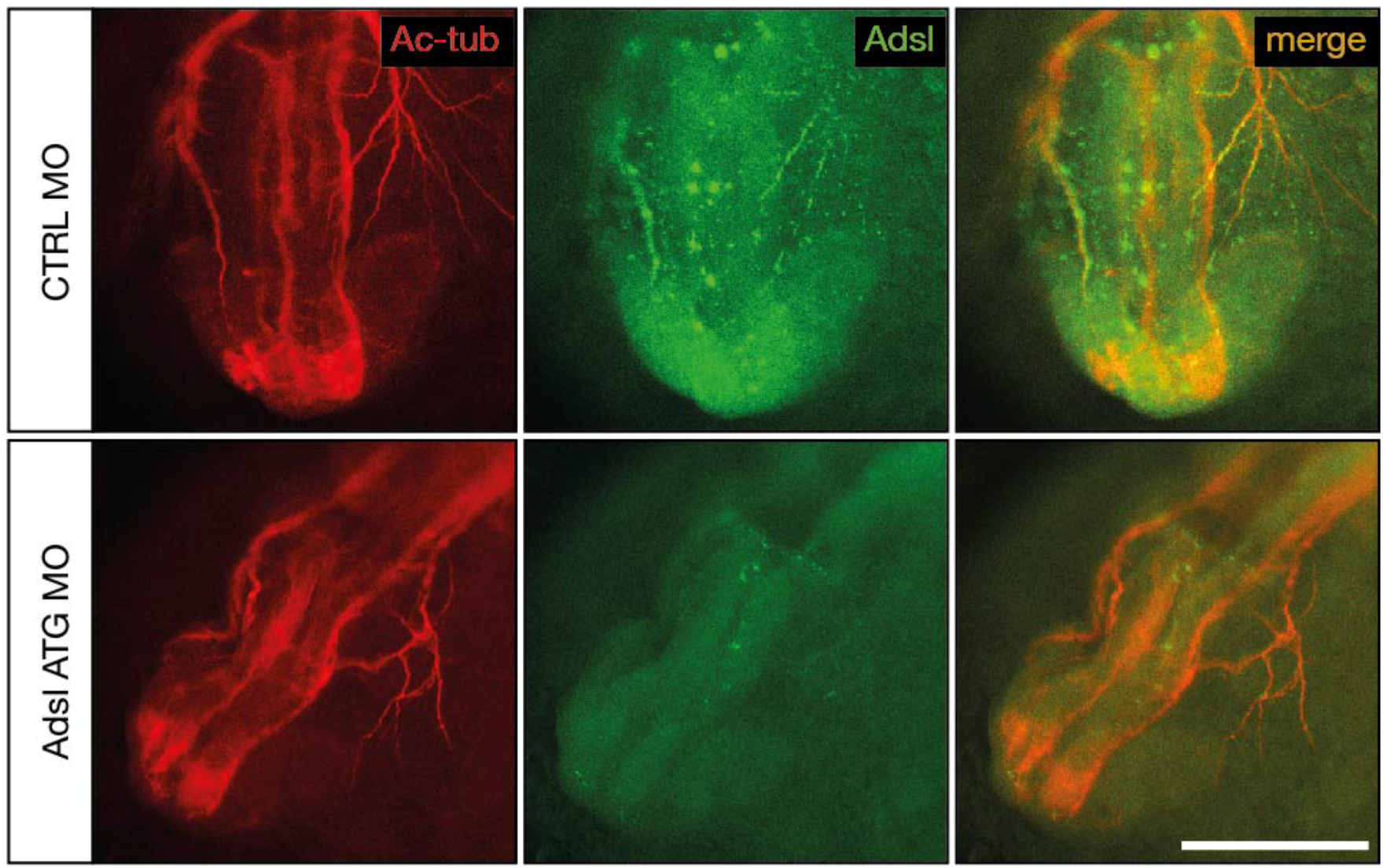
Test of knockdown efficiency. Whole mount antibody staining for acetylated tubulin (red) to visualize neurons and Adsl (green) in 24 hpf zebrafish embryos. Injection of ATG morpholino resulted in very weak expression of Adsl along axons. Images show anterior views of zebrafish heads. Scale bar=200 μm.

### Adsl depletion impairs ciliogenesis in zebrafish

As the observed phenotypes were potentially indicative of defects in cilium function, we examined heart looping by staining for cardiac myosin light chain 2 *(cmcl2)* mRNA. Adsl depleted embryos showed higher frequencies of defects, including inverse looping and to a lesser extent no loops (Figure 7A). Inverse heart looping may be indicative of laterality impairment (situs inversus) that can arise due to ciliary defects. To corroborate this possibility, we examined liver placement by staining for angiopoietin-like 3 *(angptl3).* A significant increase in inverse liver placement was observed in Adsl depleted embryos compared to controls, supporting a general defect in laterality (Figure 7B). To further investigate the laterality defects, we examined left-right asymmetry at the 20-somite stage, staining for the mRNA of the left lateral plate mesoderm marker *southpaw (spaw).* Consistent with the altered distribution of *cmcl2* and *angptl3,* asymmetric *spaw* mRNA localization was changed in about 40% of Adsl depleted embryos. Most of these embryos showed symmetric patterning and a smaller fraction no or only weak staining. The correct asymmetric distribution of *spaw* mRNA could be largely restored by expression of mRNA encoding zebrafish Adsl (Figure 7C).

**Figure 7.**
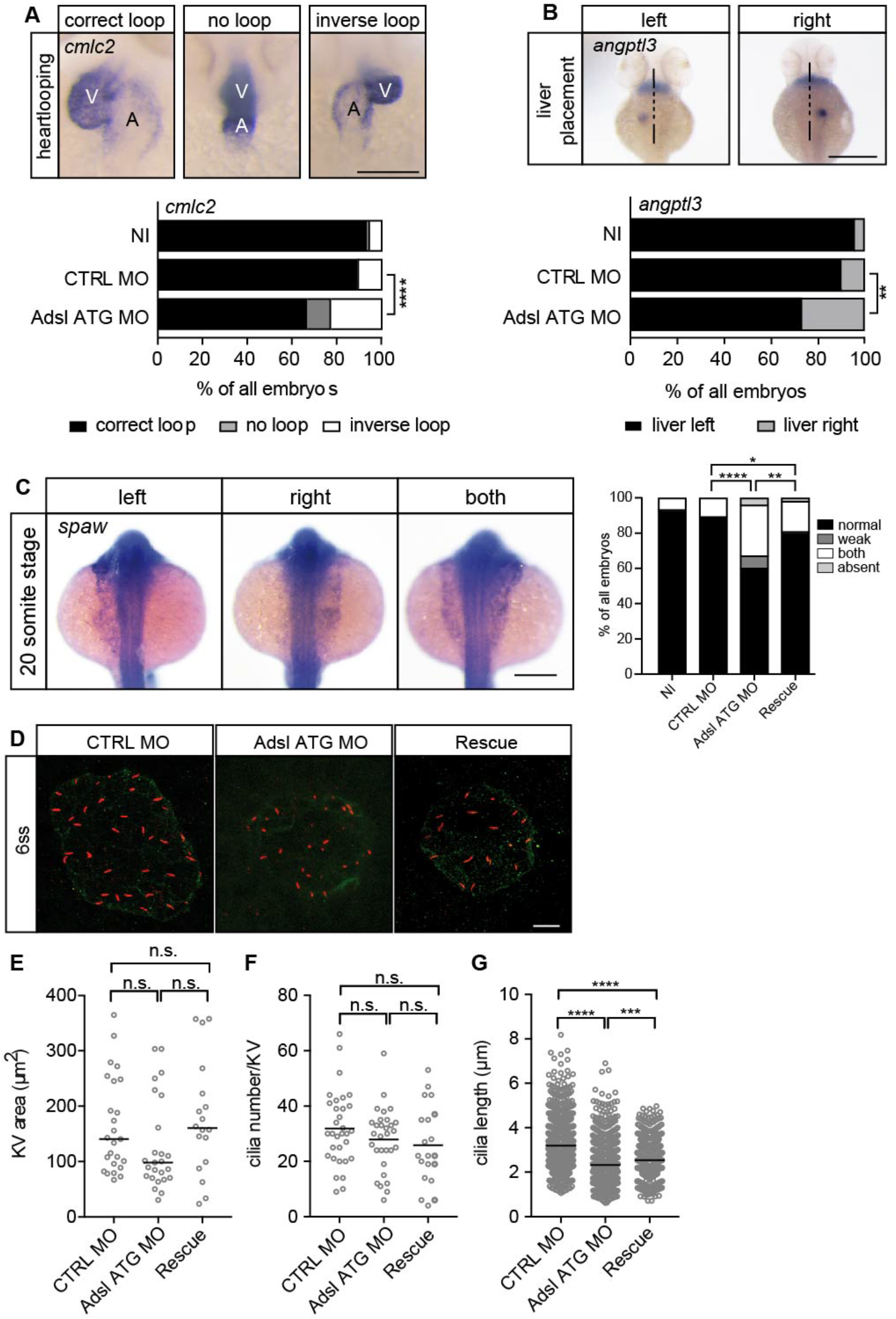
Impaired LR asymmetry and cilium formation in the organ of laterality. **A.** At 48 hpf ventricle (V) of the two-chambered zebrafish heart is placed left and above the atrium (A). Adsl depleted embryos more frequently develop inversely looped hearts or developed unlooped hearts (no loop) (as scored by whole mount *in situ* hybridization for *cardiac myosin light chain 2 (cmlc2)).* N=6 experiments with a total of 266 embryos (NI), 176 embryos (CTRL MO), 188 embryos (Adsl ATG MO). Scale bar=100μm. **B.** Whole mount *in situ* hybridization for *angiopoietin-like 3 (angptl3)* to assess liver placement in 48 hpf embryos. Dorsal view. Scale bar=200μm. 185 NI, 121 CTRL MO and 99 Adsl ATG MO embryos. (A, B) Two-tailed Fisher’s exact test; **p<0.0015, ****p<0.0001. **C.** Whole mount *in situ* hybridization for the left lateral plate mesoderm marker *southpaw (spaw)* at 20 somite stage (ss). S*paw* is normally expressed in the left lateral plate mesoderm. When LR asymmetry is disturbed, *spaw* can be detected on the right side or on both sides. Aberrant expression of *spaw* in Adsl morphants. Co-injection of RNA encoding zebrafish Adsl restores proper *spaw* expression. Two-tailed Fisher’s exact test; *p=0.0451, **p =0.0016, ****p<0.0001. Results from 5 experiments with 121 embryos (NI), 142 (CTRL MO), 128 (Adsl ATG MO) and 105 (Rescue) are shown. Scale bar=200μm. **D.** Confocal z-stacks of the Kupffer’s vesicle (KV) of 6 somite stage (ss) embryos. Cilia are stained red (acetylated tubulin), while apical cell borders were stained for PKCζ (green). Scale bar=10μm. **E.** No significant changes in the size of the KV upon Adsl depletion. n= 25 (CTRL MO), 25 (Adsl ATG MO) and 18 embryos (rescue with zebrafish *adsl* RNA). Each circle is one embryo, line indicates median. Kruskal-Wallis test with Dunn’s correction. p-values: CTRL MO vs. Adsl ATG MO: 0.2582, CTRL MO vs. Rescue: >0.9999, Adsl ATG MO vs. Rescue: 0.1684. **F.** No significant changes in the number of cilia per KV. n=32 (CTRL MO), 30 (Adsl ATG MO) and 20 embryos (rescue with zebrafish *adsl* RNA). Each circle is one embryo, lines show means. One-way ANOVA with Sidak’s multiple comparison test. p= 0.5538 (CTRL MO vs. Adsl ATG MO), 0.2844 (CTRL MO vs. Rescue), 0.9225 (Adsl ATG MO vs. Rescue). **G.** Shorter cilia in Adsl morphants can be partially elongated by coinjection of RNA encoding zebrafish Adsl. n= 960 cilia (CTRL MO), 798 (Adsl ATG MO), 540 (Rescue). Kruskal-Wallis test with Dunn’s correction, lines indicate medians; ***p=0.0008.

To test if impaired laterality may involve ciliary defects, we examined the Kupffer’s vesicle (KV, organ of laterality). While KV area and cilia number were not significantly affected by Adsl depletion, cilia length was reduced in Adsl ATG MO treated embryos, a phenotype that was partially rescued by co-injection of RNA encoding zebrafish Adsl (Figure 7D-G). These data, in combination with additional phenotypes, including laterality defects and hydrocephalus, support the role of ADSL in promoting proper cilia formation or function *in vivo*.

### MTX treatment rescues neurogenesis in Adsl depleted zebrafish

As ciliogenesis defects were metabolite dependent in human cells, we examined the effects of inhibiting purine synthesis at steps prior to ADSL in the DNPS pathway during zebrafish development. We quantified the effects of Adsl depletion on differentiating neuronal cells by staining for the marker Elavl3/4. Similar to what we observed in the chicken neural tube, depletion of *adsl* caused a significant reduction in Elavl3/4 positive cells that could be rescued by the co-injection of RNA encoding zebrafish Adsl (Figure 8A). We next treated control and Adsl depleted embryos with MTX to attenuate the DNPS pathway upstream of ADSL and reduce SAICAR production. Treatment with MTX completely rescued the reduction in Elavl3/4 positive cells in the neural tube, indicating that this was not a result of impaired DNPS *per se,* but a consequence of intermediate metabolite accumulation (Figure 8B). Similar results were observed with a second morpholino targeting (Figure S7). We next examined the effect of MTX treatment on Sox2 positive neural progenitors in the developing forebrain (Figure 8C, upper right panel). Depletion of Adsl reduced the number of Sox2 positive cells and this was rescued by cotreatment with MTX (Figure 8C). These data indicate a specific role of SAICAr accumulation in the neural progenitor defects associated with Adsl depletion.

**Figure 8.**
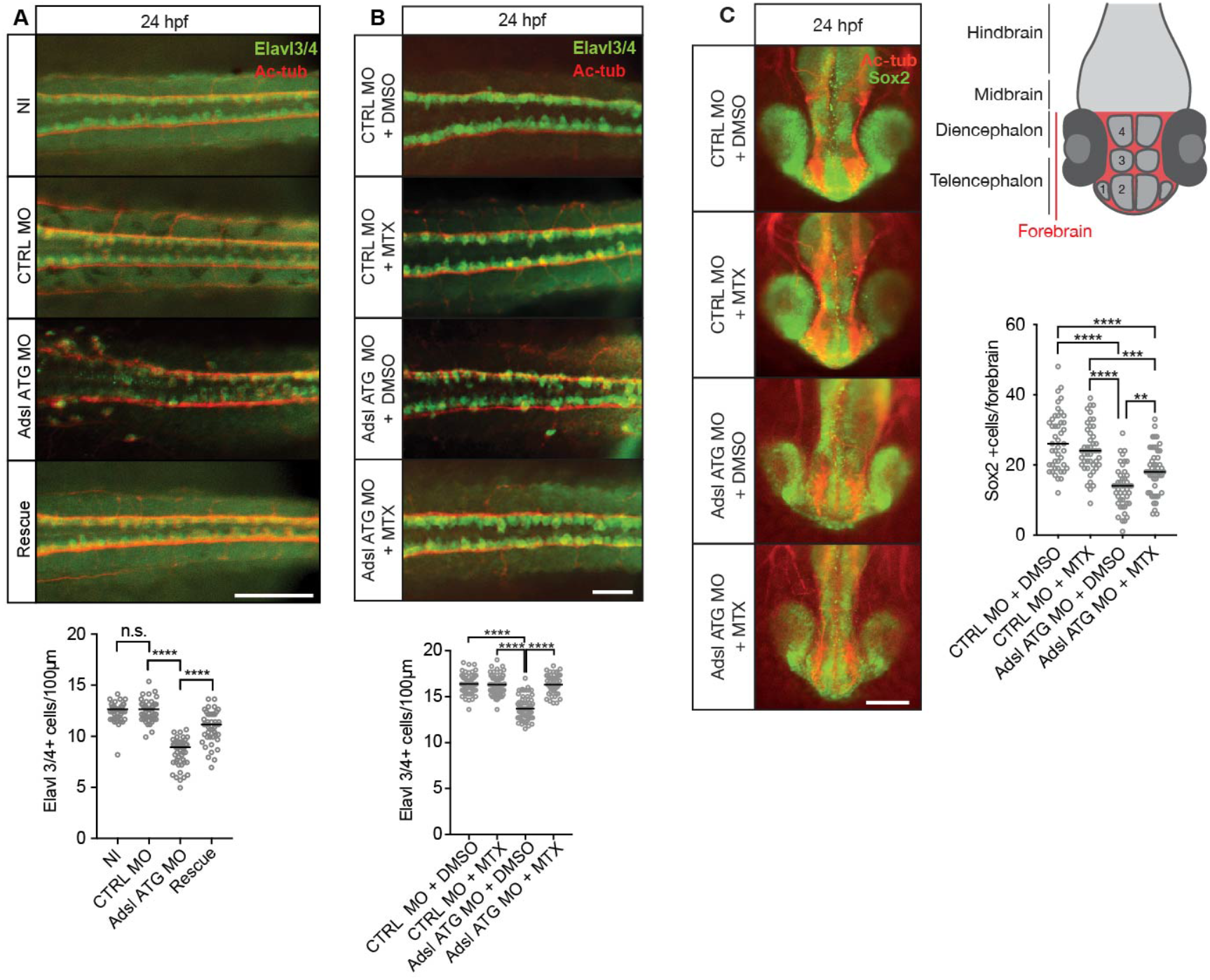
Adsl depletion reduces neuronal lineage cell numbers that can be rescued by Methotrexate treatment. **A.** Immunofluorescence whole-mount microscopy of neural tubes of 24 hpf zebrafish embryos (dorsal view) stained for acetylated tubulin (axons, red) and Elavl3/4 (green). Fewer Elavl3/4 positive cells in Adsl depleted embryos, that can be rescued by co-injection with RNA encoding zebrafish Adsl (Rescue). Graph shows Elavl3/4 counts of individual embryos, line indicates median. 3 experiments with 45 embryos (NI), 45 (CTRL MO), 45 (Adsl ATG MO), 45 (Rescue). Kruskal-Wallis test with Dunn’s correction. ns p>0.9999, ****p<0.0001. Scale bar=100μm. **B.** Methotrexate (MTX) treatment rescues Elavl3/4 positive cell numbers. Staining of the neural tube (dorsal view) of 24 hpf zebrafish embryos for acetylated tubulin (red) and or Elavl3/4 (green). Adsl morphants show fewer Elavl3/4 positive cells, which could be rescued by treatment with 100 μM MTX. 5 experiments with 69 (CTRL MO), 75 (CTRL MO + MTX), 63 (Adsl ATG MO) and 58 (Adsl ATG MO + MTX) embryos. One-way ANOVA with sidak’s multiple comparison. ns p>0.9999, ****p<0.0001. Scale bar=50μm. **C.** Forebrains of 24 hpf zebrafish embryos (left panels) stained for acetylated tubulin (red) and Sox2 positive neural progenitors (green), anterior view. Scale bar=200μm. Schematic of the developing brain of zebrafish embryos adapted from *(19)*, top right panel. The forebrain (red) is composed of the telencephalon with the olfactory bulb (1), the pallium (2), the optic recess region (3) and the diencephalon with the hypothalamus (4). Quantification of phenotypes (bottom right panel). Adsl morphants show fewer neural progenitor cells in the forebrain, that can partially be rescued with 100 μM MTX from tailbud stage on. Data were analyzed using one-way ANOVA with Sidak’s multiple comparison. Dashes show medians. Experiments with 45 embryos (CTRL MO + DMSO), 45 embryos (CTRL MO + MTX), 45 embryos (Adsl ATG MO + DMSO), 47 embryos (Adsl ATG MO + MTX). If not shown in the graph all other comparisons are not significant.

**Supplemental Figure S7.**
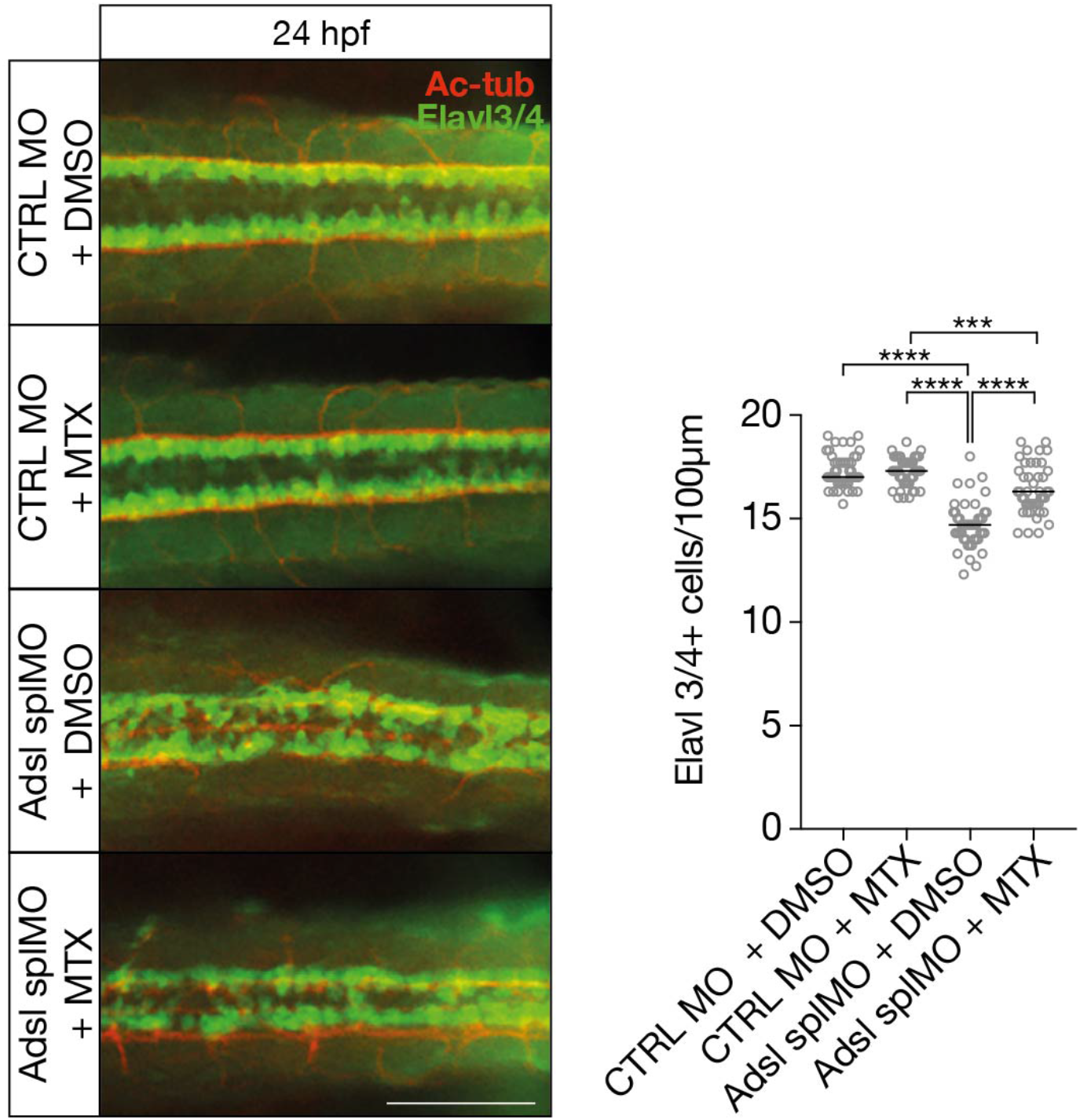
Dorsal view of part of the neural tube of 24 hpf zebrafish embryos (left panels). Acetylated tubulin (Ac-tub) is shown in red. Neural cells are stained for Elavl3+4 (green). Adsl morphants (splMO) show fewer neuronal cells, which can be partially rescued by treatment with 100 μM Methotrexate from tailbud stage on. Data were analyzed by using one-way ANOVA with Sidak’s multiple comparison. Dashes show median. Experiments with 45 embryos (CTRL MO + DMSO), 45 embryos (CTRL MO + MTX), 45 embryos (Adsl splMO + DMSO), 45 embryos (Adsl splMO + MTX). If not shown in graph, all other comparisons are not significant. Scale bar=100μm

## Discussion

Despite a detailed understanding of the enzymology of the DNPS pathway, the specific cell and organismal effects underlying the complex etiology of ADSLD remain unclear. ADSLD-linked clinical phenotypes were proposed to results from toxic accumulation of metabolites such as SAICAr, but the cellular defects that may result from increased SAICAr levels and the contribution of purine deficiency, if any, have not been investigated.

Our results uncovered multiple phenotypes in both human RPE-1 cells, as well as developing chicken and zebrafish, that can be rescued by distinct interventions. DNA damage signaling was suppressed by nucleoside supplementation, suggesting that this was caused by purine deficiency that we could readily detect in ADSL depleted human cells. In contrast, defects in primary ciliogenesis were rescued by PAICS inhibitor or MTX and phenocopied by SAICAr administration, indicating that they resulted specifically from SAICAr accumulation. In ADSL depleted RPE-1 cells, we could also detect p53 activation and defects in cell cycle progression, in the absence of cell death or senescence. These phenotypes were insensitive to nucleoside supplementation or SAICAr modulation, indicating the involvement of additional pathways.

Microcephaly, which is present in a subset of more severe ADSLD patients, was observed in zebrafish embryos following Adsl depletion. Similarly, ADSL depletion in chicken embryos led to a reduction in neural tube size. Together, the results suggest that they are potentially valuable models for understanding the etiology of the disease(*3*). DNA damage, p53 activation and defects in cilia function, that we observed following ADSL depletion, have all been implicated in neurodevelopmental disorders associated with microcephaly(*20*). One prominent example is Seckel Syndrome. In patients with mutations in *ATR,* Seckel Syndrome is caused by progenitor cell death due to replication stress and DNA damage(*21, 22*). This is accompanied by extensive p53 activation, but co-deletion of p53 exacerbated the cellular and organismal phenotypes, indicating a protective effect of p53 induction. In addition, mutations in centrosomal proteins, such as CENPJ/SAS4/CPAP or CEP63, have also been implicated in Seckel Syndrome(*23, 24*). In contrast to mice expressing hypomorphic *Atr*, progenitor loss in mice with CEP63 or CENPJ/SAS4 deficiency resulted from mitotic delays and the activation of the USP28-53BP1-dependent mitotic surveillance pathway(*25–31*). In this case the phenotype was completely rescued by p53 co-deletion, revealing p53-dependent cell death as main driver of the phenotype. Despite the phenotypic similarities at the cellular level, we did not detect increased cell death as result of ADSL deficiency, indicating that the reduced cellularity following ADSL depletion is mechanistically distinct from Seckel Syndrome.

We propose that the reduction in brain size resulting from ADSL depletion is largely due to the impaired cell cycle progression observed in SOX2 positive progenitors. This is supported by the overall reduction in SOX2 positive cells that we observed in both chicken and zebrafish embryos after ADSL depletion. As a result, differentiated ELAVL3/4 positive cell numbers were also reduced in both systems. Cell cycle delay is further supported by the observation that contrary to overall Sox2 positive cell number, the number of transfected, Sox2-positive cells in chicken embryos was increased relative to controls. In agreement with our findings, nutrient restriction was shown to arrest the proliferation of neural progenitors of *Xenopus larvae* and zebrafish reversibly in G2, suggesting that most of the cells were quiescent(*32*). While arrest did not require mTOR signaling, nutrient dependent cell cycle reentry was mTOR dependent. Purine deficiency was shown to inhibit the mTORC1 pathway, which regulates protein synthesis in response to nutrient availability(*14*). Inactivation of mTORC1 results in microcephaly in mouse models, but similar to Seckel Syndrome, this was attributed mainly to increased levels of cell death(*33*). Thus, while we cannot rule out a role for dysregulated mTOR in the phenotypes associated ADSLD, it is unlikely a major contributor.

In zebrafish, we could largely rescue neural progenitor loss by treatment with MTX, indicating that this was likely not the result of defects in overall purine synthesis, but due primarily to the accumulation of specific metabolites, most likely SAICAr. While there are currently no treatments for ADSLD, current clinical trials (NCT03776656) are examining the efficacy of the HPRT substrate analog allopurinol in an attempt to reduce the production of SAICAr and S-Ado in ADSLD patients. While our experiments focused on embryonic effects on the neural progenitor population, SAICAr accumulation may also affect primary cilia in the post-natal brain. The molecular targets of SAICAR remain largely unclear but its accumulation has been linked to activation of PKM2, and other kinases, in the context of glucose deficiency in cancer (*10–12*). Recent work has also implicated ADSL in the activation of MYC, that plays a major role in controlling metabolism and proliferation in cancer cells(*34*). Considering that ciliogenesis is a highly regulated process and tightly coordinated with the cell cycle, modulation of central signaling pathways that control energetics and proliferation may be one way by which SAICAr could affect cilia. Alternatively, SAICAr could impinge on more direct regulators of ciliogenesis. To address this future work will have to determine the cellular SAICAr interactome and identify the disease-relevant targets.

To our knowledge, this is the first demonstration of a specific purine metabolite impairing ciliogenesis. While ciliopathy-like features have not been described for the pathology of ADSLD, we observed a robust rescue of the neural progenitor population following MTX treatment, suggesting that SAICAr and its effects on cilia may be involved. Consistent with effects on cilia *in vivo,* we observed shorter cilia in the KV of zebrafish, as well as several ciliopathy related phenotypes, consistent with impairment of ciliogenesis or cilia function by SAICAr accumulation on *in vivo.* Adsl-depleted zebrafish also presented with defects in skull cartilage formation that is coordinated with brain size, in part through cilia based sonic hedgehog signaling(*35*). Primary cilia and Hedgehog signaling have well established roles in regulating multiple progenitor populations in the developing brain(*36*). However, we also note that severe defects in primary cilia function or Hedgehog signaling cause more drastic reductions in progenitor proliferation and numbers than we observed in either system, potentially consistent with the milder ciliogenesis effects observed in vivo(*37, 38*). Moreover, we have used ADSL knockdown in our experimental systems and it is currently unclear to what extent the observed ciliary defects would be recapitulated by ADSL mutations in patients and in what tissues and cell types. If ADSL deficiency would be less severe or only a subset of tissues would be affected, patients may not present classic and widespread ciliopathy features. Together, our work provides the first cell-level analysis of ADSL deficiency, identifies specific cellular defects, and ascribes these to either SAICAr accumulation or purine deprivation. Highlighting the complex etiology of ADSLD, our results add further support to the notion that SAICAr plays a key role and establish a framework for deciphering the underlying molecular mechanisms.

## Materials and Methods

### Human cells culture

Human immortalized hTERT-RPE-1 WT, TP53 knock out (kind gift from Brian Tsou), RPE-1 expressing pLenti-EGFP and pLenti-ADSL*EGFP (siRNA resistant mutant) cells were cultured in Dulbecco’s modified Eagle Medium-F12 (DMEM-F12; Thermo Fisher Scientific) supplemented with 10% (v/v) fetal bovine serum (Millipore Sigma) and 100 U ml^-1^ penicillin-streptomycin at 37°C and 5% CO2 in humidified atmosphere. For cilia experiments, silenced RPE-1 cells were serum starved for 48 hrs in OptiMEM (Thermo Fisher Scientific).

### Drugs used and concentrations

1 mg/ml SAICAr (Carbo Synth) was added to the cells for 96 hrs, to mimic ADSL depletion. 60 μM nucleosides (100X Embryomax, Merck Millipore) were added from the first silencing to the end at 1X in the culture medium. MRT00252040 (kindly provided by Simon Osborne, LifeArc, London, UK) dissolved in DMSO was used at 2 μM and MTX (Millipore Sigma) at 4 μM as described in (*14*). ATM inhibitor (KU-55933; Selleckem) was used at 5 mM for 24 hrs before fixation. Doxorubicin (Millipore Sigma) was used as positive control for senescence at 1 ug/ml for 6 days.

### siRNA transfections

RPE-1 (hTERT-RPE-1; ATCC) were transfected with 100 nM siRNAs (Millipore Sigma or Dharmacon) with Lipofectamine RNAiMAX (Thermo Fisher Scientific) in Opti-MEM (Gibco) without antibiotics for one or two rounds of 48 hrs, depending on the gene to be silenced. We used siGFP (GGCUACGUCCAGGAGCGCCGCACC) and siGL2 (CGUACGCGGAAUACUUCGA) as negative controls (siC). In this study we used a smart pool (four siRNAs) against *ADSL* (Dharmacon) or single oligos siADSL#2 5’-CAAGAUUUGCACCGACAUA-3’ (Millipore Sigma). The siRNA-resistant mutant was designed to be resistant to siADSL#2. For rescue experiments with siCP110 we used three oligos (#1 5’-GCAAAACCAGAAUACGAGAUU-3’, #2 5’-CAAGCGGACUCACUCCAUATT-3’ and #3 5’-TAGACTTATGCAGACAGATAA-3’ (Millipore Sigma) for 24 hrs.

### RNA extraction and quantitative real time-PCR

RPE-1 cells (ATCC) were seeded in a 6 well-plate, silenced for 96 hrs, washed twice in PBS and resuspended in 300 μl of Tri-Reagent (Millipore Sigma). RNA was isolated by centrifugation followed by chloroform extraction, isopropanol precipitation, washing twice in 75% ethanol and resuspended in 20 μl DEPC-treated water (Thermo Fisher Scientific). Total RNA was quantified with a Nanodrop 8000 Instrument (Thermo Fisher Scientific). 1 μg of total RNA was used for the reverse transcription reaction performed by High Capacity RNA-to-cDNA Kit (Applied Biosystems), according to the manufacturer’s recommendations, in a 2×RT buffer mix, supplemented with dNTPs, random primers and RT enzyme in a final volume of 20 μl. Quantitative real time PCR (qRT-PCR) was performed using the comparative CT method and a Step-One-Plus Real-Time PCR Instrument (Thermo Fisher). Amplification of the 16 ng of cDNA was done in triplicate with TaqMan Universal PCR Master Mix (Thermo Fisher) for *ADSL* and *GAPDH.*

### Plasmid cloning and generation of stable cell line

Human ADSL^WT^ cDNA was PCR amplified using KOD Hot start DNA polymerase (Millipore) according to manufacturer’s instructions (primers: forward containing 5’-BsiWI (ADSL-BsiWI-F-5’AAAACGTACGATGGCGGCTGGAGGCGATCAT3’) and reverse primer containing *3’-EcoR1* restriction sites *(ADSL-EcoR1-R:* 5’TTTTGAATTCCAGACATAATTCTGCTTTCA3’). PCR products were purified by using PureLink Quick Gel Extraction kit (ThermoFisher) and cloned into pCR2.1-TOPO vector (Invitrogen). Omnimax competent E. coli cells were transformed with the pCR2.1-TOPO clones and colonies were selected in carbenicillin. Constructs were then sequenced with primers for the TOPO vector (T7 Promoter-F and M13-R). By using the restriction enzymes AscI and Not1-HF (New England Biolabs), *ADSL* was cut from the TOPO vector and after gel purification, it was ligated into the MCS-BioID2-HA vector, a gift from Kyle Roux (Addgene plasmid #74224; http://n2t.net/addgene:74224; RRID:Addgene_74224)(*39*) with Quick ligation kit (BioLabs). Omnimax competent *E.coli* cells were transformed and selected with carbenicillin. The construct was confirmed by restriction digestion and sequencing (Macrogen). The human ADSLD patient mutation R426H was generated using the QuikChange mutagenesis kit (Thermo Fisher) with the following primers: ADSL^R426H^, FW, 5’-AGGCATCAACCTGGATATGCTCTATGAGGTCATTG-3’ and RV, 5’-CAATGACCTCATAGAGCATATCCAGGTTGATGCCT-3’. For complementation experiments, we cloned ADSL^WT^ cDNA and the siRNA resistant mutant into the pLenti-CMV-eGFP-BLAST (659-1) plasmid, a gift from Eric Campeau & Paul Kaufman (Addgene plasmid #17445; http://n2t.net/addgene:17445; RRID:Addgene_17445)(*40*), using primers containing *XhoI* and *EcoRI* overhangs (ADSL-*XhoI* FW 5’-AAAACTCGAGCGATGGCGGCTGGAGGCGATCAT-3’ and ADSL-*EcoRI*-RV 5’-TTTTGAATTCCAGACATAATTCTGCTTCA-3’). The siRNA resistant mutant was produced by introducing 5 different silent mutations using the QuikChange mutagenesis kit (Thermo Fisher) with the following primers: forward, 5’-GGTTTGCCAGGAGGCGTAGGTCTTTGCAAATTGTGTGCACTGATGCCCCCA-3’. And reverse 5’-CCAAACGGTCCTCCGCATCCAGAAACGTTTAACACACGTGACTACGGGGGT-3’. Constructs were checked by sequencing (Macrogen) and expression was checked by western blot and immunofluorescence. For virus preparation: 6 ×10^6^ AD293 cells were plated in 15 cm culture dishes, and transfected with 20 μg pLenti-CMV-EGFP empty and pLenti-CMV-ADSL*-EGFP, 2 μg REV, 6 μg RSV-RRE and 2 μg VSV-G plasmids with 160 μl PEI pH 7.0 (Polyscience Euro) and 150 mM NaCl. After 48 hrs the medium containing the viruses was cleared with a 0.45 mm filter (Millipore) and added to the target cells. Three days after the infection, cells were selected with blasticidin (Invitrogen) for 7 days.

### Immunofluorescence (human cells)

Silenced RPE-1 cells were seeded on 18 mm round coverslips after 96 hrs of silencing and fixed accordingly with the antibody requirements, with 4% PFA for 10 or 30 min, followed by 0.1% Triton-PBS for 5 min and stored in 100% EtOH. Cells were incubated with the blocking solution of 3% bovine serum albumin (Millipore Sigma) in PBT for 30 min. Primary antibodies (listed below) were diluted in the same blocking solution and incubated for 1 hr at RT. After three washes, cells were incubated with Alexa Fluor-conjugated 594 and 488 secondary antibodies (Thermo Fisher) at 1:400 dilution for 1 hr at RT. DAPI was used to visualize the DNA. Slides were imaged using Orca AG camera (Hamamatsu) on a Leica DMI6000B microscope equipped with 1.4 100X oil immersion objective. AF6000 software (Leica) was used for image acquisition. Image processing and quantification was performed with ImageJ software. Intensities were measured in images acquired with the same exposure settings and subtracting the background for each image.

### Antibodies

Staining of human cells was performed with the following primary antibodies: a-ADSL (Millipore Sigma, rabbit, 1:100 IF, 1:1000 western), a-ARL13B (Santa Cruz Biotechnology, mouse monoclonal C5, 1:100), PCNT (Novus Biologicals, rabbit, 1:400), a-p53 (Cell Signaling, mouse monoclonal 1C12, 1:100), a-RPA (Calbiochem, mouse monoclonal Ab-3, 1:100) a-53BP1 (Novus Biologicals, rabbit, 1:400), a-pSer139-H2A.X (Santa Cruz Biotechnology, rabbit, 1:100), a-Actin (Millipore Sigma, mouse monoclonal AC-40, 1:1500), a-Vimentin (Abcam, rabbit, 1:100), a-CK20 (DaKo, mouse, 1:200), a-Centrobin (a kind gift from Ciaran Morrison, mouse, 1:500 (*41*)), a-Centrin (EMD Millipore, mouse, 1:1,000), a-CP110 (a kind gift from Andrew Holland, rabbit, 1:1000).

Staining of chicken tissues was performed with the following primary antibodies: a-ELAVL3/4 (Molecular Probes Molecular Probes, mouse, 1:500), a-β-TubulinIII-Tuj1 (Covance, mouse, 1:1000), Pax6 (DSHB, mouse, 1:250), SOX2 (Invitrogen, rabbit, 1:500), pH3S10 (Millipore, rabbit, 1:500), Cleaved-Caspase-3 (Millipore, rabbit, 1:500). Staining of zebrafish tissues was performed with the following primary antiobodies: a-ELAVL3/4 (GeneTex, rabbit, 1:1000), a-acetylated-alpha-tubulin (Sant Cruz Biotechnology, mouse monoclonal 6-11B-1, 1:1000), a-SOX2 (Abcam, rabbit, 1:1000), a-γH2AX (GeneTex, rabbit, 1:400), a-ADSL (Millipore Sigma, rabbit, 1:200) and a-PKCζ (Sant Cruz Biotechnology, rabbit, 1:500).

### Cell proliferation and cell death

150,000 RPE-1 cells were plated in 6-well plates and silenced with control or siADSL oligos (Millipore Sigma) for 72 hrs, when they were counted and plated again in the same amount for the second round of silencing. After 3 days cells were counted as second timepoint (144 hrs, 6 days) and seeded for a third timepoint (9 days). Cells were cultured in the presence of serum for all the experiment. The ΔPDL (difference in population doubling levels) was obtained by using the formula: log(N1/N0)/log2, where N1 is the number of cells at the timepoint we collected them and N0 is the initial number of cells plated(*42*). For detecting cell death, cells in suspension were collected in the growth medium and the attached ones were trypsinized and resuspended in complete medium to block trypsin activity. Cells were then mixed in 0.4% trypan blue solution (Gibco). The number of blue-positive cells and total cell number was quantified at the microscope.

### Cell extracts and western blotting

RPE-1 cells were seeded in a 6-well plate and after 96 hrs of silencing they were trypsinized, washed once in PBS and resuspended in a 2X SDS lysis buffer (2X SDS lysis buffer contained 4% SDS, 20% glycerol, 120 mM Tris/HCl pH 6.8, 1x protease (Roche) and phosphatase inhibitors (Millipore Sigma)). Protein concentration was quantified using the *DC* Protein Assay (Bio-Rad), and proteins separated by SDS-PAGE and transferred to 0.2 μm nitrocellulose membrane (Amersham Protran) or 0.45 μm PVDF membrane (Millipore Sigma) depending on the molecular weight. Membranes were blocked in 5% milk in PBT (PBS containing 0.2% Tween-20) for 30 min and then incubated with primary antibodies for 1 hr at RT. After three washes in PBS containing Tween-20 0.02%, membranes were incubated with secondary antibodies conjugated to HRP and protein bands were visualized by ECL-Plus (Millipore Sigma).

### Senescence-associated (SA) β-galactosidase assay

RPE-1 were silenced for 96 hrs with siControl and siADSL#2, then fixed in ice-cold X-gal fixative solution (containing 4% formaldehyde, 0.5% gluteraldehyde, 0.1 M sodium phosphate buffer pH 7.2) for 4 minutes. After two washes in PBS, X-gal (Roche) was diluted 1:100 at a final concentration of 1 mg/ml in X-gal solution (containing 5 mM K_3_Fe(CN)_6_, 5 mM K_4_Fe(CN)_6_, 2 mM MgCl_2_ in PBS). Incubation was performed at 37°C for 8 hrs in the dark. Two washes in PBS were performed before taking the images. Doxorubicin was used as a positive control.

### Statistical analysis

*In vitro* data were analyzed with an unpaired two-sided *t*-test when two samples were compared, while one-way ANOVA was used to compare more than two samples in the same graph (GraphPad Prism 6.0, GraphPad Software Inc.). Values of p<0.05 were considered statistically significant (*p<0.05; **p<0.01; ***p<0.001; ****p<0.0001). Two or more independent experiments were performed for each condition and this is indicated in individual figure legends.

### Cloning (fish)

To generate a template for the generation of an antisense *in-situ* probe, a 921 bp fragment of the *Danio rerio adsl* open reading frame was cloned into pCRII via TOPO TA cloning (Invitrogen). To have a template for the generation of capped mRNA using the AmpliCap SP6 High Yield Message Maker Kit (Cellscript) the whole open reading frame of zebrafish *adsl* was cloned with a N-terminal Flag-tag into pCS2+ using EcoRI and XhoI.

### Immunofluorescence (fish)

Zebrafish embryos were fixed with 4% buffered paraformaldehyde at the indicated stages. Antibody staining was performed as described(*43*) using the primary antibodies previously described (see also antibodies section above) and detected with Alexa-Fluor-labelled secondary antibodies (1:1000, Molecular Probes).

### Statistical analysis (fish)

The number of fertilized eggs per clutch determined the size of experimental groups with clutches having been randomly and equally divided into treatment groups. No additional statistical methods have been applied to pre-determine sample size. All zebrafish experiments were done at least three times with eggs from different mating tanks or different mating days. Embryo numbers are given in the legends. All statistical analyses were performed with GraphPad Prism 7 and 8, respectively. Data were tested for normality and analyzed accordingly by parametric or non-parametric tests. Graphs display, if not indicated otherwise, individual datapoints and medians in case of non-parametric datasets. An a level of <0.5 was considered significant.

### Zebrafish maintenance and manipulation

Zebrafish were maintained in a 14 hrs light and 10 hrs dark cycle in a standardized, water recycling housing system (Tecniplast) with automatic monitoring and adjustments of pH, conductivity and temperature. Fertilized eggs were generated by natural matings of the wild-type strains EK or AB. Eggs were incubated at 28.5 °C and allowed to develop until the desired stages. In order to achieve Adsl knockdown, a translation blocking antisense morpholino oligonucleotide (Adsl ATG MO) (5’-TCCCTCCATGCCTGCAGCGGTTAAA) was used or a MO which targets the exon-intron boundary at exon 4 of Adsl (Adsl SplMO) (5’-CCAACTGTGGGAGAGAGCGACTGTA). A standard control MO was also used in all experiments. MOs (GeneTools Inc) were injected at the 1-2 cell stage directly into the yolk. In addition, non-injected wild-type embryos served as internal control for clutch quality. For pharmacological manipulation zebrafish embryos were immersed in embryo water containing 1% DMSO or 1% DMSO and 100 μM methotrexate (MTX; Cayman Chemicals) from 10 until 24 hrs post fertilization (hpf) or 50 μM nucleosides. All zebrafish maintenance and procedures have been approved by the Veterinary Care Unit at Ulm University and University of Tübingen, respectively and the animal welfare commissioner of the regional board for scientific animal experiments in Tübingen, Germany. Zebrafish experiments were performed according to the European Union Directive 86/609/EEC for the protection of animals used for experimental and other scientific purposes.

### *In situ* hybridization (ISH)

Zebrafish were fixed over night at 4°C at the indicated stages using 4% buffered paraformaldehyde, dehydrated with a gradual methanol series and stored at −20 °C until further use. For ISH embryos were rehydrated in a methanol series containing PBST (PBS containing 0.1% Tween-20) and processed according to standard protocols(*44*). Genes of interest were detected using DIG-labeled *in situ* probes, which were *in vitro* transcribed from linearized plasmids carrying fragments of the gene of interest: *adsl* (Genbank no.199899.2), *angiopoietin-like 3 (angptl3,* Genbank no. AF379604). The probes against *cardiac myosin light chain 2 (cmcl2)* and *spaw* have been described before(*45*).

### Analysis of cartilage formation

4 days post fertilization (dpf) old zebrafish embryos were fixed for 2 hours at RT using 4% buffered paraformaldehyde. After rinsing with PBS, embryos were washed for 10 min with 50% EtOH in PBS, before the staining solution (0,02% Alcian blue (Millipore Sigma), 70% EtOH, 50mM MgCl_2_) was added and the embryos were incubated o/n at RT. On the next day, embryos were rinsed with H_2_O and subsequently bleached for 20 min at RT with opened lid of the reaction tube (bleaching solution: 1,5% H_2_O_2_ in 1% KOH). A clearing series was performed (30 min 20% glycerol/0,25% KOH, 2h 50% Glycerol/ 0,1% KOH). Stained embryos were stored at 4 °C in 50% Glycerol/ 0,1% KOH.

### Measurements of cilia and neural progenitors/differentiated cell populations

To count neural progenitors, anterior views of 24 hpf embryos were taken using a fluorescent whole mount microscope. The number of Sox2 positive cells within the forebrain was determined. To count differentiated neural cells, dorsal views of embryos were captured by fluorescent whole mount microscopy and the number of ELAVL3/4-positive cells per 100 μm were counted. γH2AX positive cells were counted over a distance of 300 μm in the neural tube. Cilia were counted and measured after acquiring confocal z-stacks of flat-mounted tails of 6 somite stage (ss) embryos. The Simple Neurite Tracer in Fiji was used to trace and measure cilia through the whole z-stack. Image J was also used to trace and measure the outline of the KVs.

### Microscopy of zebrafish embryos

Live zebrafish embryos and those processed by ISH or for cartilage staining were imaged using a M125 whole-mount microscope equipped with a Leica IC80 HD camera. Zebrafish embryos undergoing immunofluorescence stainings were assessed with a M205 FCA and a DFC 9000 GT sCMOS camera. Confocal z-stacks were acquired on a TCS SP5II with LAS AF software (All microscopes and software: Leica).

### Chick embryo *in ovo* electroporation

Eggs from White-Leghorn chickens were incubated at 37.5°C in an atmosphere of 45% humidity and the embryos were staged according to Hamburger and Hamilton(*46*). Chick embryos were electroporated with column purified plasmid DNA (3 μg/μl for shRNAs) in H2O containing Fast Green (0.5 μg/μl). Briefly, plasmid DNA was injected into the lumen of HH12 neural tubes, electrodes were placed on either side of the neural tube and electroporation was carried out by applying five 50 ms square pulses using an Intracel Dual Pulse (TSS10) electroporator set at 25 V. Transfected embryos were allowed to develop to the specific stages and then dissected under a fluorescence dissection microscope.

### DNA constructs

shRNAs were generated using pSHIN plasmid (a GFP expressing evolution of pSUPER): shCONTROL sequence (CCGGTCTCGACGGTCGAGT) and shADSL sequence (GAGCTGGACAGATTAGTGA). The knockdown efficiency of shRNAs was assessed by RT-qPCR in electroporated chicken embryonic fibroblast cultures(*47*).

### Immunostaining and EdU incorporation in chicken embryos

Embryos were fixed overnight at 4°C in 4% paraformaldehyde and immunostaining was performed on vibratome sections (60 μm) following standard procedures. After washing in PBS-0.1% Triton X-100, the sections were incubated overnight with the appropriate primary antibodies diluted in a solution of PBS-0.1% Triton supplemented with 10% bovine serum albumin. After washing in PBS-0.1% Triton, sections were incubated for 2 hr at room temperature with the appropriate Alexa conjugated secondary antibodies diluted in a solution of PBS-0.1% Triton supplemented with 10% bovine serum albumin. After staining, the sections were mounted and examined on a Leica SP5 or a Zeiss Lsm 780 multiphoton microscope. For EdU incorporation, 200 ul of EdU solution (1mM) was added on the vitelline membrane of each embryo 2 h before fixation in 4% paraformaldehyde. EdU was detected in sections using the Click-iT EdU imaging kit (Invitrogen).

### Fluorescence Associated Cell Sorting (FACS)

HH-12 chicken embryos were electroporated with shCONTROL or shADSL plasmids and 48 hpe, a single cell suspension was obtained by digestion for 10-15 min with Trypsin-EDTA (Millipore Sigma) and labeled with Hoechst and a-ELAVL3/4 antibody used with Alexa647-conjugaded anti-mouse secondary antibody. Alexa647, Hoechst and GFP fluorescence was determined by FACSAria Fusion cytometer (BD Biosciences), and the data were analyzed with FlowJo software (Tree Star) and Multicycle software (Phoenix Flow Systems; cell cycle profile analysis).

### Quantitative fluorescence image analysis

Quantification of Cleaved-Caspase-3 immunofluorescence intensity was done using ImageJ software. Tuj1+ and Tuj1-areas on the electroporated side and the respective areas on the non-electroporated side were delimitated by polygonal selection, and the mean intensity of Cleaved-Caspase-3 immunofluorescence was quantified as mean gray values. At least three different images were used to calculate the mean value per embryo. Each mean value was normalized to the mean value obtained for the respective nonelectroporated area of the same embryo.

## Author contributions

I.D. performed all experiments involving cultured cells, J.G. and M.B. conducted zebrafish experiments, A.H. conducted chicken embryo experiments and S.J. and O.Y. performed metabolomic analysis. I.D, J.G, M.B, A.H, C.B, S.P, M.P, J.L, and T.H.S. analyzed data and prepared figures, C.B., J.L. and T.H.S. conceived the project, M.P., J.L. and T.H.S. obtained funding and supervised trainees, I.D, J.L, and T.H.S. designed experiments and I.D, J.L, and T.H.S. wrote the manuscript with editorial contributions from all authors.

## Acknowledgements

Thanks to members of the Lüders, Roig and Stracker labs for input, A. Riera for chemistry advice, C. Morrison for Centrobin antibody, A. Holland for CP110 antibody, B. Tsou for p53-deficient RPE1 cells, D. Zafra for assistance, C. Donow and S. Burczyk for excellent help with zebrafish maintenance and LifeArc for supplying MRT00252040. I.D. was funded by the European Union’s Horizon 2020 research and innovation programme under the Marie Skłodowska-Curie grant agreement No. 754510, T.H.S. and J.L. were funded by the Ministry of Science, Innovation and Universities (MCIU; PGC2018-095616-B-I00 to T.H.S and PGC2018-099562-B-I00 to J.L.), the 2017 SGR 1089 (AGAUR), FEDER, the Centres of Excellence Severo Ochoa award and the CERCA Programme. T.H.S. was supported by the NIH Intramural Research Program, National Cancer Institute Center for Cancer Research. M.P. was funded by grants from the Deutsche Forschungsgemeinschaft (DFG PH144/4-1 and PH144/6-1).

